# A Quantum Dot Biomimetic for SARS-CoV-2 to Interrogate Dysregulation of the Neurovascular Unit Relevant to Brain Inflammation

**DOI:** 10.1101/2022.04.20.488933

**Authors:** Wesley Chiang, Angela Stout, Francine Yanchik-Slade, Herman Li, Bradley Nilsson, Harris Gelbard, Todd Krauss

**Affiliations:** Department of Biochemistry and Biophysics, University of Rochester Medical Center, Rochester, New York 14642, United States; Center for Neurotherapeutics Discovery and Department of Neurology, University of Rochester Medical Center, Rochester, New York 14642, United States; Department of Chemistry, University of Rochester, Rochester, New York 14627, United States; Departments of Pediatrics, Neuroscience, and Microbiology and Immunology, University of Rochester Medical Center, Rochester, New York 14642, United States; The Institute of Optics, University of Rochester, Rochester, New York 14627, United States

## Abstract

Despite limited evidence for competent infection and viral replication of SARS-CoV-2 in the central nervous system (CNS), neurologic dysfunction is a common post-acute medical condition reported in “recovered” COVID-19 patients. To identify a potential noninfectious route for SARS-CoV-2-mediated neurological damage, we constructed colloidal nanocrystal quantum dots linked to micelles decorated with spike protein (COVID-QDs) as a biomimetic to interrogate how blood-brain barrier (BBB) dysregulation may subsequently induce neuroinflammation in the absence of infection. In transwell co-culture of endothelial bEnd.3 monolayers and primary neuroglia, we exposed only the bEnd.3 monolayers to COVID-QDs and examined by fluorescence microscopy whether such treatment led to (i) increased inflammation and leakage across the bEnd.3 monolayers, (ii) permeability of the COVID-QDs across the monolayers, and (iii) induction of neuroinflammation in neuroglial cultures. The results of our study provide evidence of neuroinflammatory hallmarks in cultured neurons and astrocytes without direct exposure to SARS-CoV-2-like nanoparticles. Additionally, we found that pre-treatment of our co-cultures with a small-molecule, broad-spectrum inhibitor of mixed lineage and leucine rich repeat kinases led to reversal of the observed dysregulation in endothelial monolayers and resulted in neuroglial protection. The results reported here may serve to guide future studies into the potential mechanisms by which SARS-CoV-2 mediates neurologic dysfunction.

## INTRODUCTION

Despite the explosion of studies since the advent of the COVID-19 pandemic, the basis for neurologic disease that is part of “long COVID” remains unclear. While studies of SARS-CoV-2 infection of nasal tissues as a direct route for neuroinvasion have demonstrated persistent infection, the evidence for widespread infection of neurons, microglia, astrocytes (astroglia) and other neural cell types in the central nervous system (CNS) is limited (1). Indeed, recent reports have strongly suggested that acute and chronic inflammation in the periphery and CNS drives neurologic disease in long COVID (2, 3). Taken together, experimental and pathologic data suggests that CNS disease arises from dysregulation of the barrier forming cells of the neurovascular unit (NVU) (4–6). Specifically, this likely arises from compromised integrity and inflammatory status of the blood-brain barrier (BBB) either by (A) a direct interaction between SARS-CoV-2 virus particles and the endothelial cells lining the BBB or (B) an indirect mechanism through recruitment and activation of peripheral immune cells, such as neutrophils, that release pro-inflammatory factors that interact with the constituent cells of the BBB (7–9).

Key to both inquiries is understanding whether disruption and inflammation of the endothelium is sufficient to initiate neuroinflammation in the CNS, or whether virus particles traverse across the dysregulated BBB and result in subsequent physiochemical interactions with cellular constituents of the CNS to induce a neuroinflammatory state. Delineation between these two mechanisms may be determined by application of fluorescently labeled virus-like nanoparticles (VLNPs) to visualize whether SARS-CoV-2 virions are transported across the BBB into the CNS. Typically, VLNPs are constructed with hydrophobic cores loaded with conventional fluorophores or contain fluorescent protein constructs (10–12). These fluorophores, however, can be limited by smaller absorption cross sections, poor photostability, and low overall brightness that make single-molecule detection and extended biological studies challenging (13–15). These limitations can be addressed by incorporating fluorescent inorganic nanoparticles, such as quantum dots (QDs), into VLNPs (16–18). Compared to conventional fluorophores, QDs exhibit enhanced photophysical properties that result in improved sensitivity for biosensing and bioimaging applications (13, 19–21). While initial concerns regarding toxicity and general biological stability have limited the broad adoption of QDs in the biomedical field, a robust body of work has emerged that shows excellent stability and specificity of biological probes incorporating QDs (22–29).

To address how SARS-CoV-2 may initiate brain inflammation, we have constructed a SARS-CoV-2 VLNP with a QD core (COVID-QDs) to interrogate whether (i) direct, physiochemical interactions with the brain endothelium can induce NVU inflammation and (ii) if such neuroinflammation arises from direct physiochemical interactions with this SARS-CoV-2 mimicking nanoparticle. Specifically, the COVID-QDs were constructed by linking SARS-CoV-2 spike (S) proteins to polymeric phospholipid chains that formed a micellar envelope around the QDs. These COVID-QDs structurally and functionally mimicked SARS-CoV-2 virions to elicit an immunologic response in cutured cells without infection. The COVID-QDs reported here were designed to achieve such by accurately replicating the physical dimensions and average number of surface bound S proteins reported for SARS-CoV-2 virions from cryogenic electron microscopy (cryo-EM) and atomic force microscopy (AFM) measurements (17, 30, 31). Additionally, the inherent photophysical properties of the COVID-QDs allow for fluorescent readout of their localization and interaction with biomolecular targets; properties that are not inherent to most other SARS-CoV-2 virus-like nanoparticle constructs (32, 33). Here we report that COVID-QDs induce loss of endothelial tight junctions and upregulation of inflammatory molecules in an *in vitro* model of the BBB. These changes are reversible with treatment of either soluble human angiotensin-converting enzyme 2 (hACE2) or URMC-099, a broad-spectrum anti-neuroinflammatory and neuroprotective brain-penetrant mixed-lineage kinase 3 (MLK3) and leucine rich repeat kinase type 2 (LRRK2) inhibitor. Finally, we observed induction of dendritic beading, a marker of synaptodendritic neuronal injury in a co-culture model system of the NVU after S or COVID-QD treatment, which could be reversed by URMC-099. Taken together, our data support (i) COVID-QDs are a potential tool to further interrogate the neurological deficits associated with SARS-CoV-2 and (ii) that URMC-099 may have a role in adjunctive treatment to ameliorate these deficits in patients with long neurologic COVID-19.

## RESULTS & DISCUSSION

### Construction and Characterization of COVID-QDs

The COVID-QDs were constructed in a two-step process as outlined in Fig. 1A and detailed in the SI Appendix (Materials and Methods). CdSe/CdS core-shell QDs were synthesized as detailed in the SI Appendix (Materials and Methods) and then encapsulated by a polymerized phospholipid micelle, following by conjugation of S ligands to the QD-micelle surface. The micellar envelope is formed by ligands that we synthesized. Its constituents are a phospholipid (phosphatidylethanolamine; PE) polymer (polyethylene glycol; PEG) chain attached to a reactive group (bis(2-methylphenyl)sulfone; bis-sulfone) (SI Appendix, Fig. S1). The bis-sulfone group we appended to our polymerized phospholipid ligand takes advantage of the poly-histidine tags, commonly used to purify recombinant proteins, as a site of conjugation that does not perturb the tertiary structure of the protein of interest. Specifically, the bis-sulfone undergoes an efficient sequential Michael addition-elimination reaction, or bis-alkylation, selective for adjacent imidazole functional groups in poly-histidine chains (34, 35). Thus, after formation of the QD-micelle, this bis-alkylation mechanism allowed efficient, irreversible conjugation of multiple S proteins to form COVID-QD constructs.

**Figure 1.**
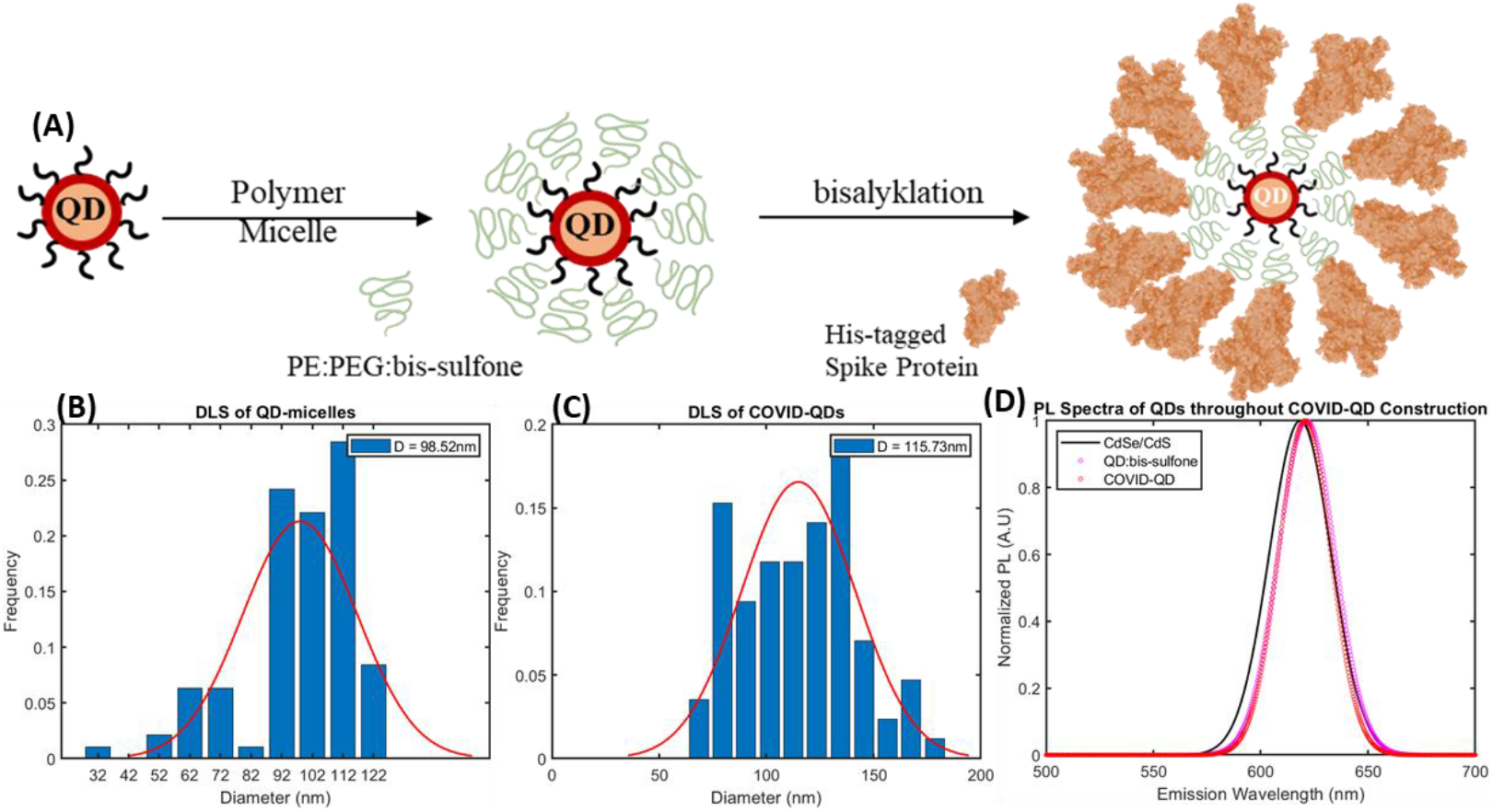
Two-step construction of COVID-QDs and corresponding size and photoluminescence analysis of constructs after each step. **(A)** CdSe/CdS QDs are synthesized and encapsulated in a polymeric micelle formed from PE:PEG:bis-sulfone ligands and then conjugated with excess ligands of S. DLS size analysis of **(B)** QD-micelles (***d_avg_*** ≈ **98*nm***) and **(C)** COVID-QDs (***d_avg_*** ≈ **115*nm***) validating size mimicry and expected increase in size corresponding to distributed S conjugation across micellar surface. **(D)** Normalized PL spectra of CdSe/CdS QDs, QD-micelles, and COVID-QDs to confirm reaction scheme does not shift the PL emission spectrum (***λ_em_*** = **620*nm***).

A micelle encapsulation protocol was selected for the ability to produce a micelle population within the size range of 80 nm-140 nm commonly reported for SARS-CoV-2 virus particles from cryo-EM and AFM studies (30, 31). After synthesis, the QDs had a diameter of 9 nm that was increased to a diameter of 98 nm after micellation and size-selective purification determined by transmission electron microscopy (TEM) (SI Appendix Fig. S3) and dynamic light scattering (DLS) measurements, respectively (Fig. 1B). After conjugation of S proteins, the COVID-QDs had a final diameter of 115 nm (Fig. 1C) that is well within the desired size range; this size increase is also proportional to having S conjugation evenly distributed across the micellar surface. The successful functionalization of COVID-QDs with multiple S proteins was confirmed by eluting the unconjugated S protein fraction after termination of the reaction and measuring absorbance at 280 nm (A280) of the eluted fraction on a NanoDrop Spectrophotometer. The A280 measurement was corrected for molecular weight and estimated extinction coefficient of S protein to calculate an approximate concentration. This value was used to determine an 87.5% ± 1.2% conjugation efficiency for the reaction (Table 1). This represents an average of 18 S molecules/COVID-QD, within the distribution of 24 ± 9 S per virus particle previously reported (36). The encapsulation and conjugation reactions result in no significant attenuation or shifting of QD photoluminescence (PL) emission (Fig. 1D).

**Table 1.** Parameters for A280 calculation to determine fraction of spike protein in eluted unbound fraction

|  |  |  |
| --- | --- | --- |
| MW | $\epsilon_{calc}$ | %*Spike*<sub>unbound</sub> |
| 135kDa | 166,860 | 12.5 ± 1.2 |

### COVID-QDs Dysregulate bEnd.3 Monolayers in a Spike Specific Interaction

The functional mimicry of the COVID-QDs was assessed in an *in vitro* model system of the BBB derived from a murine brain endothelial cell line (bEnd.3) cultured on transwell membranes. bEnd.3 monolayers responded to COVID-QDs similarly to treatment with soluble S as assessed by changes in barrier integrity and expression of inflammatory markers. Changes in barrier integrity were determined by transendothelial electrical resistance (TEER), shown in Fig. 2I, complemented by immunocytochemical analysis (Figs. 2A-D) of changes in the localization of the tight junction protein claudin 5 (CLDN-5), as shown in Fig. 2J. Compared to the untreated (sham) negative control group, both COVID-QD and soluble S treatments resulted in statistically significant decreases in TEER (p<0.05, one-way ANOVA + Holm-Sidak post hoc), indicative of decreased barrier integrity, consistent with previous observations of cultured monolayers of brain endothelial cells exposed to soluble S (6). The observed change in monolayer permeability likely arises from altered localization of junctional proteins, as indicated by the formation of tight junction (TJ) clusters (Figs. 2B-D) that are consistent with a measured decrease in the fraction of membrane-localized CLDN-5 in COVID-QD and S treatment groups (Fig. 2J). This observed change in TJ localization leading to decreased TEER is also consistent with the data from our positive control treatment with a tight junction-disrupting peptide (TJDP) derived from the first extracellular loop of Claudin-1 (37). Membrane-associated CLDN-5 was defined as CLDN-5 fluorescence colocalizing with expression of platelet endothelial cell adhesion molecule 1 (PECAM-1), which is constitutively expressed in cells of the vascular compartment and regulates vascular integrity and immune cell trafficking (38, 39). Additionally, previous reports observed induction of a pro-inflammatory state in endothelial cultures exposed to soluble S (6). Thus, to further validate the functional mimicry of COVID-QDs, we performed immunocytochemical analysis (Fig. 2K) of changes in expression of vascular adhesion molecule 1 (VCAM-1), which regulates inflammation-associated adhesion to endothelial cells and plays a role in inflammatory immune cell transendothelial migration across the BBB (40, 41). Fig. 2E-H photomicrographs depict low basal VCAM-1 expression in the sham control group, but significantly upregulated (p < 0.05, one-way ANOVA + Holm-Sidak post hoc) expression in soluble S and COVID-QD treated monolayers. Upregulation of VCAM-1 expression is not observed in TJDP-treated monolayers, suggesting that COVID-QDs and S protein initiate inflammation as well as loss of tight junctions. While our observations are likely due to cytoskeletal remodeling of junctional contacts, we performed a resazurin assay as an index of an intact mitochondrial respiratory chain in viable endothelial cells to confirm that these functional phenotypic changes are not simply a cytotoxic effect from either COVID-QDs or S protein. As shown in SI Appendix Fig. 4, we did not observe any significant difference in the viability of bEnd.3 cells treatment groups in response to our TJDP, Spike, or COVID-QD – nor from any of the other treatment groups involved later in this study compared to our sham negative control group.

**Figure 2.**
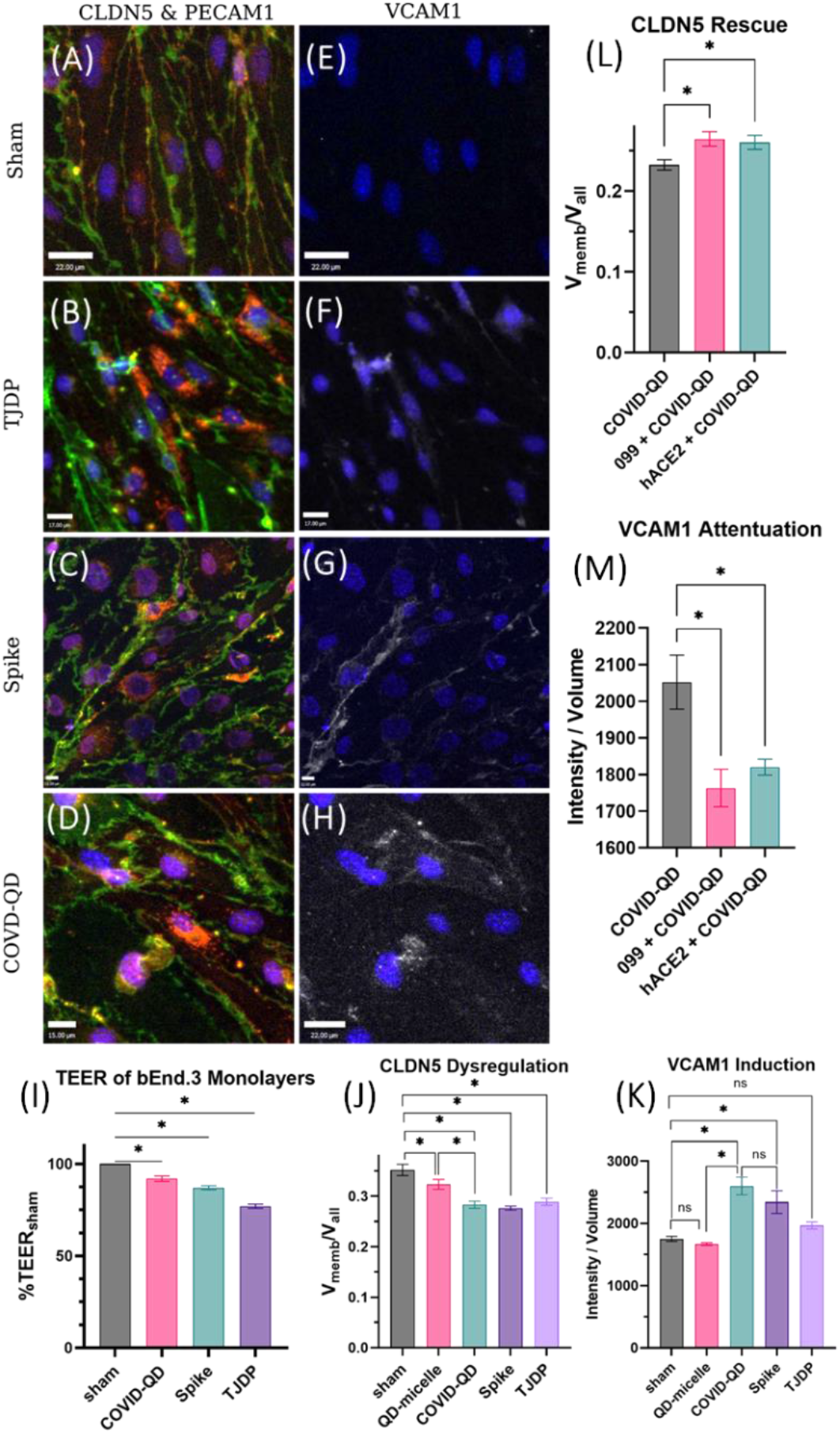
Evaluation of immunomodulation of bEnd.3 monolayers cultured on transwell inserts in response to no treatment (sham, **A & E**, scale = 22 um)), a positive control junction disrupting peptide (TJDP, **B & F**, scale = 15 um), COVID-QDs (**C & G**, scale = 15 um), and Spike protein (**D & H**, scale = 15 & 22 um). Fluorescent micrographs are immunolabeled against nuclei (DAPI, dark blue), PECAM-1 (green), tight junction protein (claudin-5, red), and VCAM-1 (white). Changes in barrier integrity are measured by a decrease in measured resistance compared to sham (**I**, *p<0.05) and are attributed to a decrease of the fraction of membrane localized claudin-5 (**J**, *p<0.05). VCAM-1 was stained in **E-H,** as represented by a white colored feature, to determine changes in inflammatory state in the bEnd.3 monolayers (**K**, *p<0.05). Unfunctionalized QD-micelles lacking Spike protein were used as an additional control to confirm COVID-QD acts in a Spike specific manner; these did not induce a change in CLDN5 localization nor VCAM-1 expression compared to sham. Health of bEnd.3 monolayers is rescued upon treatment with either soluble hACE2 or URMC-099, resulting in rescue of fraction of membrane-localized CLDN5 (**L**, *p<0.05) and attenuation of VCAM1 upregulation (**M**, *p<0.05). Statistical analyses shown are mean ± SEM (standard error of measurement) and were computed using n = 5 wells over two passages.

Lastly, to test whether the immunomodulatory effects of the COVID-QDs on the bEnd.3 monolayers is a S-specific interaction, we treated monolayers with unfunctionalized QD-micelles lacking S protein and observed no significant dysregulation of CLDN-5 localization nor induction of VCAM-1 expression (Fig. 2J & K, SI Appendix Fig. S5). This suggests that our observations are not simply due to exposure to an exogenous nanoparticle or any unreacted bis-sulfone groups on the COVID-QD surface. Additionally, we co-incubated bEnd.3 monolayers with COVID-QDs with soluble hACE2 to act as a decoy target for S proteins on the COVID-QD surface (42–45). As shown in Fig. 2L & 2M and SI Appendix Fig. S5, hACE2 co-treatment resulted in an increased fraction of membrane-localized CLDN5 and decreased VCAM-1 expression compared to monolayers treated with only COVID-QDs (p<0.05, one-way ANOVA + Holm-Sidak post hoc). These observations collectively support that COVID-QD interactions with bEnd.3 cell monolayers are S protein-specific.

### COVID-QD Mediated BBB Damage Results in Neuroinflammation of Neuroglial Cultures

The dysregulated barrier integrity and pro-inflammatory state we observed in the bEnd.3 monolayers are likely to negatively impact neuronal health and activity. Thus, to test this hypothesis, we added neuroglial co-cultures as shown in SI Appendix Fig. 6. Specifically, bEnd.3 monolayers were cultured on transwell cell culture inserts, which were then assembled above primary cultured rat hippocampal neurons and astrocytes on coverslips. The bEnd.3 monolayers act as a model BBB and segregate the basolateral (upper chamber) media from that of the apical (lower chamber) media that is conditioned with soluble factors released by the neuroglial culture. Collectively, this co-culture system partially recapitulates the static (i.e., without arterial or venous flow) aspects of the NVU. We applied the same treatment conditions described in the previous section to the basolateral media compartment and elicited similar responses in endothelial monolayers. The inserts and coverslips were separately fixed and stained for immunocytochemical analysis.

TEER and immunocytochemical analyses were performed on the bEnd.3 monolayers to confirm similar dysregulation of endothelial cells in response to the various treatments before examining the corresponding neuroglial co-cultures (Fig. 3). The leakage across the monolayers and potential activation of pro-inflammatory signaling events was expected to induce neuroinflammation in the mixed neuroglial cultures (46–48). We fixed and stained the cultures with antibodies to microtubule associated protein 2 (MAP-2), postsynaptic density protein 95 (PSD-95), and glial fibrillary acidic protein (GFAP) to identify hallmarks of neuroinflammation such as dendritic beading, synaptic pruning, and astrogliosis, respectively (49–53). We noted dendritic beading in MAP2-labeled neurons in all three treatment groups compared to the sham treatment (Fig. 3B, G, L, Q). Based on our previous work, we generated a semi-quantitative parameter for bead frequency, defined as the average number of identified dendritic beads per neuron, and corroborated that induction of beading was significant (p < 0.05, one-way ANOVA + Holm-Sidak post-hoc) in all three treatments (Fig. 3T) (51). Along with these focal swellings in dendrites, we also found a reorganization of dendritic spines as represented by a marked decrease (p < 0.05, one-way ANOVA + Holm-Sidak post-hoc) in overall PSD-95 labeling in the treatment groups compared to the sham control group (Figs. 3D, I, N, S, V). The dual observation of these focal swellings, or varicosities, and loss of postsynaptic densities are indicative of synaptodendritic injury in response to a dysregulated endothelial barrier from the S protein, COVID-QD, and TJDP treatments (51, 54).

**Figure 3.**
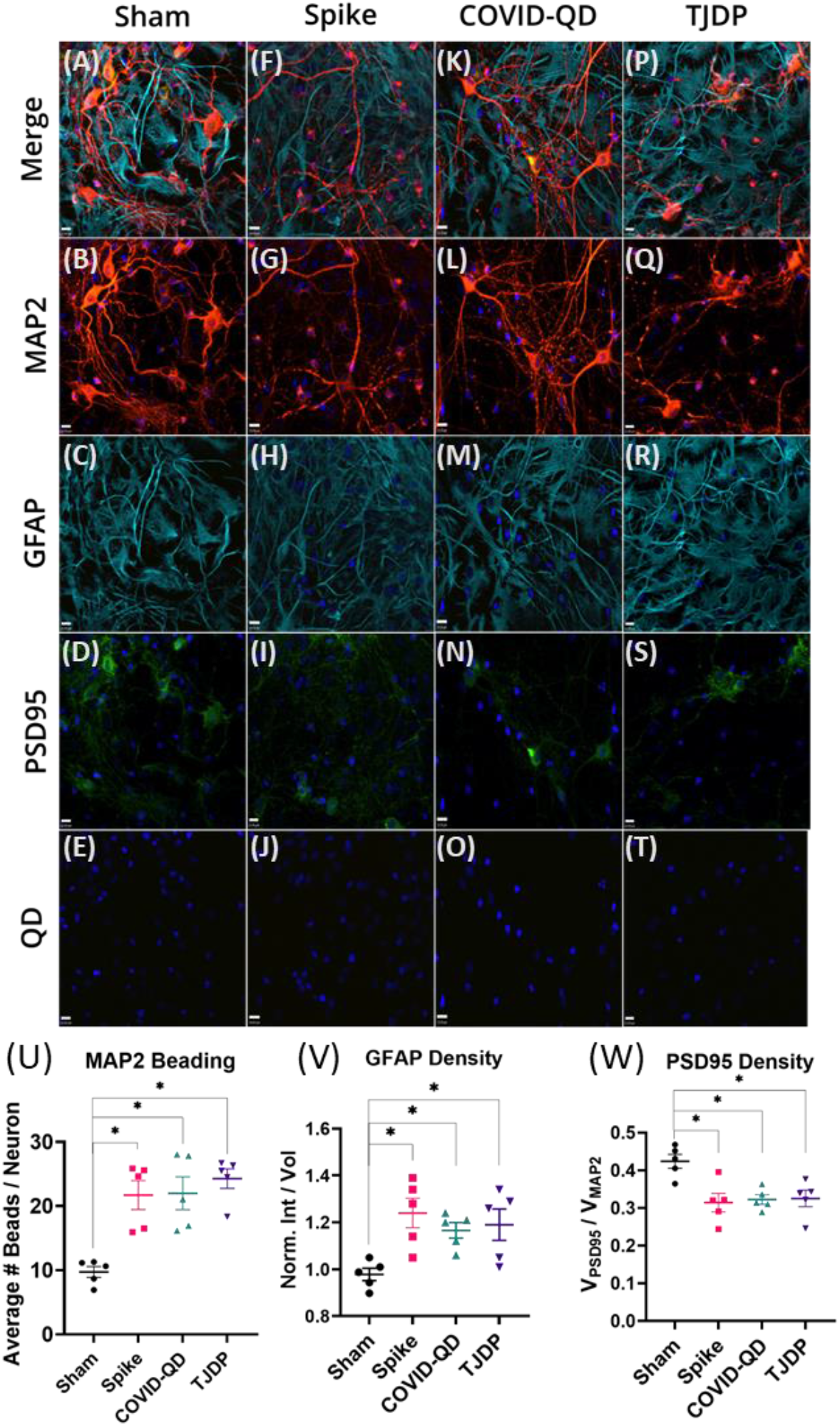
Changes in neuroinflammatory state of neuroglial culture in response to different treatment groups applied to bEnd.3 monolayers. (**A-T**) Immunofluorescent labeling of neuronal markers (red MAP2 and green PSD95), astroglia marker (cyan GFAP), and QD fluorescence (yellow puncta). Formation of puncta along dendritic processes of neurons present in MAP2 staining (**G, L, Q**) are indicative of a pro-inflammatory state in neurons and were found to be increased in treatment groups that dysregulate the bEnd.3 monolayers in comparison to the sham treatments (**U**). Changes in astrocytic state indicated by changes in GFAP expression (**V**). Expression of PSD95 lowered in co-cultures exposed to treatment groups (**W**). No observation of QD fluorescence in any of the treatment groups (**E, J, O, T**), and particularly the COVID-QD group (**O**), suggest that dysregulation of bEnd.3 monolayers does not involve potential permeability of SARS-CoV-2-like nanoparticles in out model system. DAPI stained nuclei shown in all panels. Scale bar = 15um. Plots shown are mean ± SEM and n = 5 cultures pooled from two trials.

The loss of dendritic spines represented by decreased postsynaptic densities may simply be associated with neuronal cell death resultant from permanent dendritic beading (54). However, previous reports have proposed astrocytes as potential targets in the CNS through S protein binding to alternative proteins such as the membrane-bound coreceptor neuropilin-1 to tyrosine kinase receptor for both vascular endothelial growth factor and semaphorin (55–57). Alternatively, bEnd.3 monolayers activated in response to S protein or COVID-QD treatment, may result in upregulation of complement C3 secretion that results in reactive astrogliosis (58, 59). The activation of astrocytes may contribute to synaptic elimination through multiple EGF like domains 10 (MEGF10) and MERTK, a member of the MER/AXL/TYRO3 receptor kinase family, phagocytic pathways (53, 60).Thus, our data of decreased PSD-95 expression (Fig. 4Q) may indirectly reflect astrocyte-induced synaptic pruning. Because astroglial activation in our model may also reflect brain inflammation, we examined changes in GFAP density as a representative hallmark for astrogliosis (52). From our immunofluorescent micrographs, we did find significant evidence (p < 0.05, one-way ANOVA + Holm-Sidak post-hoc) for increased GFAP density (Fig. 3U), defined by total GFAP staining intensity normalized to total stained GFAP volume, in the S protein and COVID-QD treatment groups (Fig. 3H, M) compared to the untreated group (Fig. 3C). Interestingly, while the TJDP treatment group did not induce inflammation in bEnd.3 monolayers, as indicated by VCAM-1 expression, we found a significant increase in GFAP density, indicating potential astrogliosis in this treatment group. Since the TJDP is derived from the first extracellular loop of CLDN-1, which can be expressed by reactive astrocytes in disease states such as multiple sclerosis, we presume that the observed astrocytosis is a secondary response to significantly greater leakage in the TJDP treatment groups compared to the S and COVID-QD treatment groups (61). In vivo, astrocytic end-feet comprise the glial limitans, which is the second barrier that exists between the BBB, and the perivascular space (62). Thus, TJDP exposure may disrupt normal glial limitans architecture with a compensatory response in GFAP expression. This extensive barrier breakdown and resultant leakage in our modular model NVU system in turn may explain why the TJDP reagent can induce astrogliosis, similar to observations in pathologies with extensive BBB breakdown, without upregulating VCAM-1 (60, 63).

**Figure 4.**
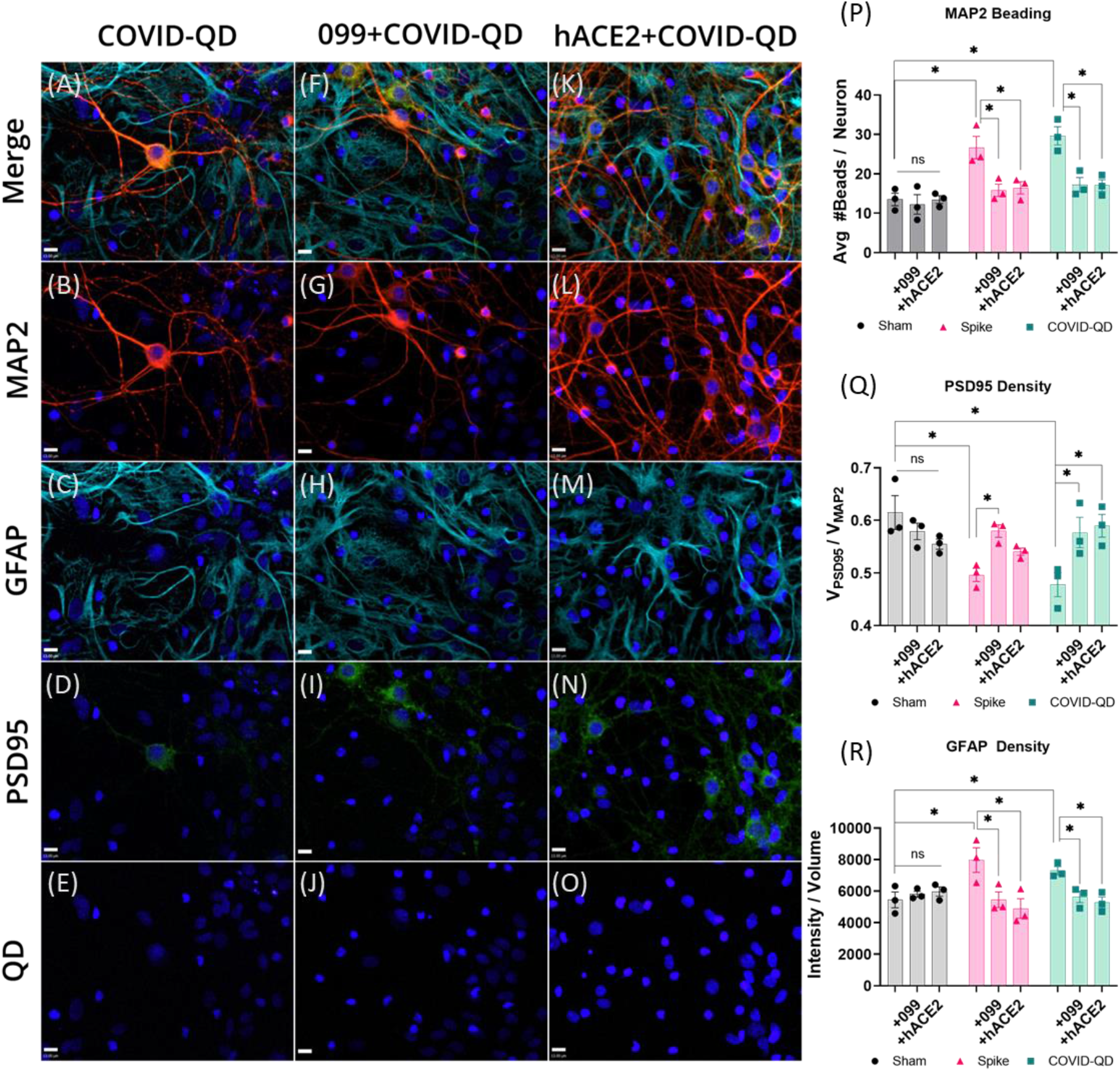
Rescue of neuronal culture in model NVU from COVID-QDs (**A-E**) with either URMC-099 pre-treatment (**F-J**) or soluble hACE2 co-treatment (**K-O**) in luminal domain of bEnd.3 monolayer. Both small-molecule treatments result in reduced dendritic beading (**P**) as indicated by the red labeled MAP2 neurons (**B, G, H**) as well as improved spatial distribution of green-labeled PSD95 (**D, I, N, Q**). A marked decrease in astrogliosis, indicated by density of cyan GFAP expression, (**C, H, M, R**) is also observed. No observable PL from QDs (**E, J, O**) in neuroglia culture. Scale bars = 15μm. Plots report mean ± SEM and n = 3 cultures.

### Neuroinflammation Does not Require Direct Physiochemical Interactions Between Neuroglia and COVID-QDs

The underlying hypothesis of this work posits that neuroinflammation of the CNS is not solely dependent on ingress of SARS-CoV-2 virions across the BBB leading to direct modulation of the constituent cells of the CNS. As a proxy for this, the COVID-QD PL was imaged using an excitation/emission optical filter pairing specific to the CdSe/CdS QDs used to construct the micelles. The resulting fluorescent micrographs (Fig. 3O) resulted in no discernable PL features indicative of COVID-QDs crossing from the luminal to abluminal domain of our model NVU. Computational thresholding of the images to find dim fluorescent puncta also resulted in no discernible identification of COVID-QDs in the neuroglial culture (Fig. 3O). Additionally, the bright fluorescent punctae visible in the QD detection channel in the corresponding bEnd.3 monolayers (SI Appendix Fig. 7) for each co-culture treatment further validates that the lack of QD fluorescence in the neuroglial cultures is not simply due to quenching of the nanoparticles. The combination of these observations lends support to the notion that COVID-QD do not directly interact with neuroglia.

### URMC-099 is a Potential Therapeutic Tool Against SARS-CoV-2 Associated Dysregulation of NVU

To further implicate endothelial barrier damage from COVID-QDs as a sufficient modulator of neuroglial health, we attempted to rescue neuroinflammation in our co-culture model using two small molecule treatments to mitigate dysregulation of the bEnd.3 monolayers. We previously showed in Figs. 2L & 2M that co-treatment with soluble hACE2 acted as a decoy target for COVID-QDs to attenuate dysregulation and inflammation of the bEnd.3 monolayers. The rescue of barrier integrity and inflammatory state of the monolayers suggested potential reduction of the observed COVID-QD associated neuroinflammatory state in our co-culture system. However, while soluble hACE2 has been theorized as a potential treatment to mitigate general SARS-CoV-2 pathogenesis, its short half-life in peripheral circulation requires recombinant fusion with Fc domains (64, 65). While our current work only examines the role of direct physiochemical interactions with the NVU, the presence of Fc domains is a concern for the indirect immunomodulatory model. The ACE2-Fc fusion constructs may activate peripheral immune cells, such as neutrophils, via binding of Fc receptors, leading to aberrant inflammation that could significantly confound the anticipated therapeutic benefits of soluble ACE2-Fc treatment (66, 67). Thus, we tested another small molecule anti-inflammatory, broad spectrum kinase inhibitor, URMC-099, that has been previously validated for NVU rescue in acute and chronic neurologic disease models (68–72). In bEnd.3 monolayers alone, pre-treatment with URMC-099 before COVID-QD treatment resulted in similar rescue of barrier health compared to soluble hACE2 co-treatment (Figs. 2L & 2M). Thus, we expected that pre-treatment of URMC-099 in our co-culture system could also result in neuroinflammatory rescue.

Following this hypothesis, we interrogated whether either hACE2 or URMC-099 rescue of bEnd.3 monolayer health in our model NVU co-culture system could robustly mitigate induction of the various neuroinflammatory hallmarks we observed in Fig. 3. Specifically, we replicated our sham, Spike, and COVID-QD treatments to the abluminal compartment of the bEnd.3 monolayers in co-culture, and additionally introduced either soluble hACE2 co-treatment or URMC-099 pre-treatment to each of the three treatment groups (Fig. 4). Correspondingly, we observed similar trends as before in the dysregulation of bEnd.3 monolayer health (SI Appendix Fig. 8) that led to induction of neuroinflammation signatures in co-cultures treated with either COVID-QDs or S protein alone (Fig. 4A-D & SI Appendix Fig. 11). Independently, neither soluble hACE2 nor URMC-099 alter the health of bEnd.3 monolayers (SI Appendix Figs. 9 & 10); thus, no markers of neuroinflammatory damage are upregulated compared to our sham control group (Fig. 4P-R). However, in line with our expectations, soluble hACE2 co-treatment or URMC-099 pre-treatment with either S protein or COVID-QDs resulted in similar rescue of bEnd.3 monolayer health (SI Appendix Figs. 9 & 10) and correspondingly reduced markers of neuroinflammatory damage (Fig. 4F-I, 4K-N & SI Appendix Fig. 11). Performing the same semi-quantitative analyses as before, we found robust reduction of dendritic beading (Fig. 4P) complemented by substantial rescue of the spatial distribution of post-synaptic densities (Fig. 4Q). We also found reduction in astrogliosis, demonstrated by reduced GFAP expression (Fig. 4R), that further suggests the potential role for astrocytic activation in mediating altered neuronal health relevant to S protein induced dysregulation of the static components of the NVU.

## CONCLUSIONS

Various studies have implicated induction of neuroinflammation as a likely mediator of altered neurologic function associated with COVID-19 neuropathophysiology (2, 73–77). This idea has gained credence from several human postmortem neuropathologic studies that have not demonstrated widespread infection of either neurons or glia as the basis for long neurologic COVID-19 (31, 76, 78). In line with these hypotheses, our data provides support for a proposed model for neurological manifestations of COVID-19 via a dysregulated BBB mediated by direct physiochemical interactions of SARS-CoV-2 virus particle mimetics with the endothelium (4, 6, 77). We show that SARS-CoV-2 mimicking nanoparticles can reduce bEnd.3 monolayer integrity by disrupting TJ formation (Fig. 2D) and inducing an inflammatory state, with increased VCAM-1 expression (Fig. 2H). This was specific for S protein interactions with the endothelium (Figs. 2J, K) that are reversible with soluble hACE2 (Figs. 2M, N). Disruption of our model BBB system results in the presentation of neuroinflammatory hallmarks in our NVU co-culture system. Specifically, neurons develop dendritic focal swelling and reduction of dendritic spines, represented by postsynaptic densities, (Figs. 3L, N, U, V). We initially thought that the modification of the neuronal processes and post-synaptic densities arises from the activation of reactive astrocytosis in response to the inflammatory state of the bEnd.3 monolayers induced by either S protein or COVID-QDs (Figs. 2G, K); this is consistent with our observations in co-culture, where S protein and COVID-QD treatments resulted in increased GFAP expression, a hallmark of astrogliosis (Fig. 3V). Interestingly, however, we also observed indications of astrogliosis in the co-culture system treated with TJDP, despite no induction of inflammation in the bEnd.3 monolayers in response to this peptide (Figs. 2F, K). Within the constraints of our current study, we posit that the small peptide TJDP may also be interacting with astrocytic end-feet due to its CLDN-1 extracellular loop like design, leading to disruption of glial limitans-like structures in our model NVU system (37, 61). Of interest for our future studies is to delineate cellular and molecular constituents that participate in inflammatory signaling pathways from the endothelium to CNS resident immune cells, as well as peripheral immune cell recruitment. This may help better explain the comparable levels of neuronal injury in all three treatment groups, with the caveat that TJDP treatment may artifactually induce astrogliosis because of the magnitude of BBB injury and subsequent exposure of neuroglia to endothelial cell media (60, 63).

Notably, our results do not include the likely involvement of peripheral immune cell mediated dysregulation of the NVU; as with sepsis associated encephalopathy, peripheral immune contributions such as granulocytes and monocyte-derived macrophages are likely to result in lasting immune activation in the CNS (7–9, 73, 79). The interaction of such cell types with SARS-CoV-2 and resultant contributions to COVID-19 neuropathophysiology are of interest for future studies. Despite lacking the contribution of this indirect mechanism of SARS-CoV-2 mediated immunomodulation of the CNS, our results show that direct physiochemical virus-host interactions at the BBB are sufficient to result in downstream neuroinflammation in CNS parenchyma. Additionally, by constructing a SARS-CoV-2 mimicking QD, we were able to assess whether neuroinflammation requires direct virus-host interactions with the constituent cells of the CNS. While we observe COVID-QD fluorescence and binding at the bEnd.3 monolayers (SI Appendix Fig. S7), no discernable fluorescence is detected in the neuroglial culture (SI Appendix Fig. S4B) despite significant induction of neuroinflammatory hallmarks, such as dendritic beading (Fig. 3L) and loss of postsynaptic densities (Fig. 3N). While this does not rule out potential interactions between the CNS and SARS-CoV-2 virus particles, our data shows that these interactions are not necessary to induce a neuroinflammatory state relevant to COVID-19 associated neurologic dysfunction. Equally salient, synaptodendritic beading represents a potential mechanism to explain synaptic dysfunction that very well may be the substrate for “brain fog”, delirium and exacerbation of pre-existing neurodegeneration, particularly in the elderly population already at risk for Parkinson’s disease and Alzheimer’s disease (51).

Lastly, in this work, we present data to support a role for a potential therapeutic, URMC-099, against SARS-CoV-2 mediated NVU injury. Therapeutically validated in various acute and chronic neuroimmune contexts including models of HIV-1 associated neurocognitive disorders as well as systemic HIV infection, URMC-099 was expected to reduce bEnd.3 barrier disruption and inflammation (80–82). We found that pre-treatment with this small molecule reduced barrier permeability, mitigated loss of membrane-localized CLDN-5, and prevented VCAM-1 induction (Figs. 2L, M). Correspondingly, rescue of the bEnd.3 monolayer health with URMC-099 also resulted in reduced induction of neuroinflammatory hallmarks, such as decreased dendritic beading, improved spatial distribution of post-synaptic densities, and attenuated astrogliosis as measured by GFAP expression. Taken together, these observations implicate direct physiochemical immunomodulation of the BBB as a potential mediator for neurologic dysfunction associated with SARS-CoV-2 neuropathophysiology and support a role for URMC-099 as adjunctive treatment to mitigate this dysregulation of the NVU. Future experiments will focus on functional readouts of altered neuroglial function associated with these data. Additionally, future transcriptomic and proteomic assays are needed to elucidate a more concrete mechanistic immunomodulatory pathway for SARS-CoV-2 mediated neuroinflammation.

## MATERIALS AND METHODS

The complete materials and methods for the experiments relevant to this work can be found the SI Appendix. For ease of following the general flow of experiments, we have included an abbreviated description of the methods used in this study.

### Construction of COVID-QDs

Synthesis of the QDs occurred on a Schlenk line under N2 with heating and stirring. Briefly, CdSe cores were synthesized from a protocol adapted from those previously reported, where Cd and Se precursors were reacted with various coordinating ligands where after a brief nucleation step at elevated temperature, the nanocrystals were annealed for up to 2 hours to achieve desired core size, as determined by absorbance and PL (15, 83). The CdSe QD cores were washed and then shelled with CdS following a protocol adapted from those previously reported, where Cd and S precursors were slowly injected via a dual-syringe pump (15, 84, 85). The final CdSe/CdS core/shell QDs were then encapsulated into self-assembled micelles formed by a polymerized phospholipid ligand with a bis-sulfone functional group (PE:PEG:bis-sulfone). The formed QD-micelles were subjected to successive rounds of filtration and size selection purification to isolate a size population most similar to native SARS-CoV-2 virions. These QD-micelles were then “decorated” with a 20x excess of His-tagged S protein via a bisalkylation reaction to form the desired COVID-QDs.

### Cell Culture and Treatment

The single culture model BBB and co-culture model NVU were constructed using an immortalized murine brain endothelial cell line (bEnd.3) for the former, as well as primary Sprague Dawley rat hippocampal cultures for the latter. The bEnd.3 cells were grown in transwell inserts to form monolayers, as determined by light microscopy (confluence > 90%) and TEER (plateauing resistance). The neuroglia were cultured on poly-D-lysine coated coverslips at embryonic day 18 and used for biological experiments between 18-21 days in vitro. To assemble the model NVU, bEnd.3 monolayers on transwell inserts were introduced to wells containing neuroglia cultured coverslips a day before treatment to allow equilibration. The cultures were then incubated for up to 18 hours with the various treatment groups before subsequent assays (TEER, cell viability, immunocytochemistry) were conducted. For ICC, cultures underwent a standard indirect immunolabeling protocol where they were fixed with 4% paraformaldehyde, permeabilized, blocked with bovine serum albumin, treated with 1° antibodies overnight at 4°C, followed with 2° antibodies for 1 hour at room temperature, before eventual mounting onto coverglass.

### Acquisition and Analysis of Immunofluorescent Micrographs

The prepared microscope slides with cells were imaged on a microscope system setup in a “grid” confocal geometry using a structured illumination element inserted into the optical path. Multicolor z-stack image sections were acquired per sample and analyzed using Volocity 3D Image Analysis Software to extra semi-quantitative measurements.

## ACKNOWLEDGEMENTS

We would like to acknowledge the expertise of Drs. Matthew Brewer and Benjamin Miller at the University of Rochester Medical Center for providing the TJDP reagent and access to their EVOM2 instrumentation to take TEER measurements. We also would like to acknowledge Dr. Steve Brochinni at the University College London for his expert insight into the bisalkylation reaction scheme relevant to PE:PEG:bis-sulfone conjugation to His-tagged S protein. We also want to acknowledge the Integrated Nanosystems Center and the Structural Biology & Biophysics facility at the University of Rochester for providing access to the TEM and DLS instrumentation, respectively. The work presented here was partially funded from the National Science Foundation (NSF; Construction and characterization of CdSe/CdS QDs, PE:PEG:bis-sulfone, COVID-QDs), National Institute of Health (NIH; Cell culture, resazurin assay, immunocytochemistry), and American Heart Association (AHA, MALDI-ToF measurements by FY). The corresponding award numbers are as follows: NSF CHE 1904847 (TDK), NSF CHE 1904528 (BLN), AHA 18CSA34020064 (BLN), NIH R21 AG074232 (HAG), NIH R01 AG057525 (HAG, supplemental URMC site award), NIH T32 GM135134 (WC), NIH T32 GM118283 (FY), NIH Office of the Director S10OD030302 (MALDI instrumentation).

## COMPETING INTEREST STATEMENT

HAG is the Chief Science Officer of Pioneura Corp, (Fairport, NY) which holds the exclusive license for URMC-099, but did not contribute either salary support or funding for this work.

## DATA AVAILABILITY

Files associated with the data reported here are accessible at OSF.io (https://osf.io/5q7ye/?view_only=3467ab2052164a75b0144c8f9b19a900).

## Notes

https://osf.io/5q7ye/?view_only=3467ab2052164a75b0144c8f9b19a900

