## Supplemental Methods and Data for "A Quantum Dot Biomimetic for SARS-CoV-2 to Interrogate Dysregulation of the Neurovascular Unit Relevant to Brain Inflammation"

### MATERIALS AND METHODS

#### Synthesis of CdSe Cores

*Reagents*: All reagents were used as purchased from the manufacturer without further purification. The following were purchased from Sigma-Alrich: trioctylephosphine (TOP; cat: 718165), selenium pellets (Se; cat: 209643), diphenylphosphine (DPP; cat: 252964), cadmium oxide (CdO; cat: 202894), trioctylphosphine oxide (TOPO; cat: 223301), 1-hexadecylamine (HDA; cat: 445312), tetradecylphosphonic acid (TDPA; cat: 736414).

*1M TOP:Se*: Prior to synthesis, 1M TOP:Se was synthesized following a protocol as previously reported (1). In a glovebox, 680mg Se pellets and 8.5mL TOP were combined in a scintillation vial and mixed at 60^o^C overnight to result in a clear solution. A small amount of secondary phosphines, 90uL DPP, was added to help promote efficient QD nucleation (2).

*Protocol*: The CdSe cores for the QDs were synthesized using a procedure adapted from those previously reported (1, 3-5). We added 820mg CdO, 16.2g TOPO, 37g HDA, and 3.2g TDPA to a 250mL 3-neck flask and heated the flask to 90^o^C under N_2_. The flask was then degassed by 3 cycles of evacuation(<100mT) and refiling with N_2_ before leaving the mixture under N_2_ and heating to 320^o^C with rapid stirring. Complexation of Cd-TDPA was visually determined by the transition from an opaque to translucent solution, upon which the flask was cooled to 260^o^C. While maintaining rapid stirring, 8.0mL 1M TOP:Se was rapidly injected and the cores were grown for 2 to 3 hours until the desired size was achieved. This was determined by taking PL measurements of aliquots every 10 minutes after the 2hr timepoint. The reaction as then quenched by removing the flask from heat and applying forced air to bring the solution to 200^o^C before submerging the flask into a water bath to rapidly cool the solution to 100^o^C. The solution was then injected with 40mL of ButOH and allowed to cool for ~1hr before performing several rounds of washing followed by two cycles of size-selective precipitation. The final pellet was resuspended in hexane and filtered through a 0.45μm syringe filter to produce the CdSe stock used for shelling.

#### Shelling of CdSe/CdS QDs

*Reagents:* All reagents were used as purchased from the manufacturer without further purification. The following were purchased from Sigma-Aldrich: oleic acid (OAc; cat: 364525), octadecylamine (ODA; cat: 74750), oleylamine (OAm; cat: O7805), octadecene (ODE; cat: O806), octanethiol (OT; cat: 471836).

*Cd-oleate Preparation*: Prior to synthesis, Cd-oleate was prepared similar to previously reported (1, 2). To a flask was added 250mg CdO, 2.6mL OAc, and 20mL ODA. The contents were degassed at room temperature, followed by heating to 270^o^C for 90 minutes under N_2_. The flask was then cooled to 150^o^C and injected with 1.3mL OAm. The Cd-oleate product was stored in a glovebox until needed.

*Protocol*: The CdS shelling procedure is a modified protocol from that previously reported (1, 4, 5), where 100nmol of the CdSe stock solution, 3mL OAm, and 3mL ODE were added to a 100mL 3-neck flask and degassed at room temperature for ~1hr under constant stirring at 800rpm. The CdSe mixture was then kept under vacuum and stirring while heated to 115^o^C for 20 minutes. The flask was then refilled with N_2_ and heated to 350^o^C at a ramp rate of 16^o^C/min. During this time the Cd and S precursors were individually loaded into two separate syringes, where 0.150mmol of Cd-oleate (~2.2mL) was diluted to 3.5mL with ODE in one syringe and 0.180mmol of OT (~0.04mL) was diluted to 3.5mL with ODE in the other. The syringes were affixed to a dual syringe pump at injected at a rate of 1.5mL/hr once the solution reached 200^o^C. The reaction was held at 350^o^C for shell growth before cooling to 200^o^C and followed with dropwise addition of 1mL OAc. The solution was left to anneal for 1hr, cooled to 75^o^C, then transferred to falcon tubes for three rounds of washing. The final pellet was resuspended in either hexane or toluene and stored in a glovebox.

#### Synthesis of DSPE-PEG_2k_-bis(2-methylphenyl)sulfone (PE:PEG:bis-sulfone)

*Reagents:* All reagents were used as provided by the manufacturers without further purification. The polymeric phospholipid was purchased from Nanocs Inc. (DSPE-PEG_2k_-NH_2_; cat: PG2-AMDS-2k). The bis(2-methylphenyl)sulfone-NHS-ester was purchased from BroadPharm (Bis-sulfone NHS Ester; cat: BP-23344). Triethylamine (TEA; cat: 15791) was purchased from Acros Organics.

*Protocol*: The synthesis of PE:PEG:bis-sulfone was carried out via a modified NHS-ester crosslinker conjugation in dichloromethane, adapted from Zhang *et al.* (6). In a 1:1 mole ratio, DSPE-PEG_2k_-NH_2_ and bis-sulfone NHS ester were added to a 25mL 2-neck flask and degassed for 1hr at room temperature. During this time a 10-mole excess of TEA was diluted in dichloromethane until an approximate pH of 8.5 was achieved using a pH strip pre-wetted with nanopure water. The flask was then refilled with N_2_ and the TEA solution was slowly injected. The reaction was left under N_2_ and gentle stirring (~300rpm) for 24 to 48 hours until the reaction was determined to be complete. The reaction progress was monitored by thin layer chromatography by using 7.5% methanol:chloroform as the eluting solvent and short-wave UV irradiation to visualize the migration of the spotted aliquots and reagents. MALDI-ToF (SI Appendix S1) and NMR were used to confirm that the desired product was formed.

*MALDI-ToF:* The 1μL of product was spotted for MALDI with 1μL of a matrix comprised of α-Cyano-4-hydroxycinnamic acid, 0.5% trifluoroacetic acid, and 0.1% NaCl dissolved in EtOH and measured at 60-80% power (Shimadzu Axima Performance).

*NMR:* For NMR, the product was dried using a rotary evaporator and redissolved in CDCl_3_ with a small aliquot of toluene added as an internal reference standard to determine a relative concentration using 1H NMR peak integration. All 1H and 13C spectra were acquired using an Avance 500 (Bruker) at 298K and the resultant peaks were manually integrated.

#### Construction of COVID-QDs

*Encapsulation of QDs in PE:PEG:bis-sulfone micelles:* An aliquot of CdSe/CdS core/shell QDs were dried under a rotary evaporator and resuspended in CHCl_3_. The QDs and PE:PEG:bis-sulfone reagents were separately aliquoted at a 1:50,000 mole ratio and briefly sonicated for 2 minutes (power level 6, VWR; model 250D) to disperse any small QD clusters or pre-formed micellar structures. The two reagents were then combined in a vial and diluted to 10x with CHCl_3_ before briefly vortex mixing for 1 minute followed by sonication for 3 minutes. The solution was then dried under a rotary evaporator. The dried gel-like film was then briefly annealed at 80^o^C in an oil bath for 10 minutes and then resuspended in ddH_2_O and stirred at 300rpm for 1-3 hours at RT. Following this, the QD-micelle solution was then passed through a size exclusion spin column (Cytiva; cat: 27513001) to remove any unencapsulated QDs, free PE:PEG:bis-sulfone ligands, or overtly large micelles. This should result in a clear solution that is then passed through a 100k MWCO Amicon Ultra Centrifugal filter (Millipore Sigma; cat: UFC810024) was used to concentrate down the QD-micelle solution and remove any QD-micelles that would be too small to replicate a natural SARS-CoV-2 virion. This product was then diluted 4x with ddH_2_O and then centrifuged at 16,000xg for 45 minutes to isolate to distinct size groups of QD-micelles. The pellet was resuspended with ddH_2_O. The eluent from the 100k MWCO filter, the supernatant, and the resuspended pellet were spotted onto a 384-well microplate and characterized with dynamic light scattering (Wyatt DynaPro Plate Reader II) and photoluminescent spectra. Photoluminescent spectras of QD, QD-micelle, and COVID-QD were imported onto MATLAB and fitted with Gaussian approximations, normalized, and plotted as shown in Fig. 1D.

*Dynamic Light Scattering (DLS):* The DynaPro system was controlled using the Dynamics software (Wyatt; v7.10) to acquire and preliminarily process the gathered spectra. Each sample was spotted in two separate wells with 10 3-second acquisitions being acquired over each well per run and each run being repeated three times. The resultant acquisitions were processed using the “Legacy” fitting algorithm on Dynamics, and then filtered for further analysis based on manual inspection of the resultant correlogram for each individual acquisition. The remaining acquisitions were then imported onto MATLAB to be plotted into a frequency normalized histogram with a Gaussian-fitted distribution overlayed as shown on Fig. 1B.

*Conjugation of Spike protein to QD-micelles:* Using the results of the dynamic light scattering analysis, the solution containing the optimal size distribution best matching native SARS-CoV-2 virions was selected. To get a rough approximation of concentration, the dilution corrected intensity of the PL from was compared to that of the QDs before micelle encapsulation. Before introducing the Spike protein, the bis-sulfone groups on the QD-micelle surface were activated by an elimination reaction to produce the mono-sulfone form of the ligand that then undergoes the bisalkylation conjugation with the poly-His tag on the Spike protein (7, 8). This was done by concentrating down the QD-micelles using a 30k MWC Amicon Ultra Centrifugal filter (Millipore Sigma; cat: UFC203024) and diluting 4x in a 50mM sodium phosphate buffer pH 7.4 + 100mM NaCl. The solution was then incubated at 37^o^C for 4-8 hours and then a 20x molar excess of Spike protein dissolved in 50mM sodium acetate buffer pH 5.6 + 35uM hydroquinone was added. This solution was mixed on a rocker for 18 hours at RT. The reaction was then quenched with 1mM sodium borohydride in 4^o^C for 90 minutes. When the solution was then filtered with a 150k MWCO centrifugal protein concentrator (Thermo Scientific; cat: 89920) to remove any unconjugated Spike protein. The concentrated COVID-QD product was then stored at 4^o^C and used within 48 hours for biological experiments. The eluent containing the free Spike protein was then concentrated down with a 30k MWCO Amicon Ultra Centrifugal filter (Millipore Sigma; cat: UFC203024) and then spotted onto a NanoDrop Spectrophotometer for an absorbance @ 280nm (A280) measurement corrected for molecular weight and estimated extinction coefficient to determine concentration.

#### Cell Culture

*bEnd.3 cell line:* The immortalized murine brain endothelial cell line (bEnd.3) was purchased from ATCC (cat: CRL-2299) and maintained according to manufacturer protocols. Specifically, the bEnd.3 cells between passage numbers 24-30 were maintained in a flask containing Dulbeco’s Modified Eagle Medium (DMEM, ThermoFisher; cat: 10567022) + 10% Fetal Bovine Serum (FBS, Atlas Biologicals; cat: F-0500-D) and passaged every 3 days with 0.25% trypsin-EDTA (ThermoFisher; cat: 25200056). For resazurin viability assays, the bEnd.3 cells were sub-cultured onto a 96-well plate (Corning; cat: 353377) at a density of 35k/well. For use in biological experiments, the bEnd.3 cells were sub-cultured onto 0.4μm pore diameter PET transwell inserts (Greiner Bio-One; cat: 662640) at a seeding density of ~85k cells/well. The formation of monolayers on these inserts were assessed daily by examining the confluency of the cells under a light microscope and taking complementary transendothelial electrical resistance (TEER) measurements (World Precision Instruments; EVOM2) after confluency appeared to be at least 80%. Monolayers of greater than 90% confluency and stagnating increases in TEER were then used in biological experiments.

*Rat hippocampal neuroglial 1^o^ culture:* Sprague Dawley (SD) rats at embryonic day 18 (E18) were used to prepare the primary neuroglial cultures for the *in vitro* model neurovascular unit setup in these experiments. The care and use for these animals were in accordance with the Guide for the Care and Use of Laboratory Animals, with protocols approved by the University Committee on Animal Resources at the University of Rochester. Hippocampi were dissected from the E18 SD rats and dissociated in 0.25% trypsin, followed by seeding of the cells at a density of 45k/well onto poly-D-lysine coated coverslips (Neuvitro; cat: GG-12-15H) in a 24-well plate. The cells were maintained in neurobasal media (ThermoFisher; cat: 21103049) supplemented with B27 with antioxidants, 1% GlutaMAX (ThermoFisher; cat: 35050061), 25μM glutamic acid, and 5% FBS. Every 3-4 days, half of the conditioned media was aspirated off and replenished with neurobasal media supplemented with B27 without antioxidants and 1% GlutaMAX. The neuroglial cultures were used in biological experiments between 18-21 days in vitro.

*Assembly of model NVU co-culture system:* Fully formed bEnd.3 monolayers on transwell inserts were introduced to wells containing neuroglial cultures on coverslips 24 hours before use in biological experiments.

#### Cell Viability Assay

Changes in cell viability in response to the various treatments used in this study was carried out using an alamarBlue (Thermo Scientific; cat# 88952) resazurin metabolism fluorescent assay. The bEnd.3 cells were plated at a cell density of ~30k/well in a 96-well plate (Sigma; cat# CLS3904) in DMEM + 10% FBS 48hrs before treatment. After the cells reached >90% confluency, as assessed by light microscopy, the cells were primed for treatment in a reduced serum condition (DMEM + 1% FBS) for 3 hours, followed by an overnight (~18hr) incubation with the selected treatments shown in SI Appendix Fig. S4. Specifically, the reagents – a sham no treatment group, 200nM URMC-099, 10nM soluble hACE2, 10nM Spike protein, 10uM TJDP, 1nM QD-micelles, 1nM COVID-QDs – were prepared in DMEM + 1% FBS and reflected the unique reagents that the bEnd.3 cultures would be exposed to. At the 15hr timepoint, the alamarBlue reagent was added to each well at a final concentration of 10% v/v. At the 18hr timepoint, a spectrophotometer plate reader ($\lambda_{ex}=550nm, \lambda_{em}=590nm$) was used to assess the degree of resazurin metabolism in each treatment group. Blank wells containing only 10% v/v alamarBlue in the reduced serum media were used to correct for baseline fluorescence. Viability measurements were conducted over two passages for a total of 7 replicates, with the measured fluorescence normalized to a sham, no treatment group from each passage to correct for passage-to-passage variability.

#### Treatment of Cell Cultures

*Dysregulation of bEnd.3 monolayers in single culture and NVU co-culture:* Prior to treatment, the cells were incubated in a reduced serum environment (DMEM + 1% FBS) for 3 hours. Following this, the abluminal domain of the bEnd.3 monolayers were exposed to a sham media only group, 10nM of Spike protein, 1nM COVID-QDs, 10uM TJDP, or 1nM of QD-micelles in DMEM + 1% FBS. The cultures were incubated in these treatments for 18hrs. Each independent treatment group was repeated for a total of 5 replicates over two separate passages of bEnd.3 monolayers as well as NVU co-cultures (two passages of bEnd.3 monolayers and 1^o^ neuroglia from separate rats).

*Small molecule rescue of bEnd.3 monolayer health:* For soluble hACE2 rescue, an equimolar concentration of soluble hACE2 was co-incubated with 10nM Spike protein in DMEM + 1% FBS for 30 minutes prior to treatment. A 10x molar excess of soluble hACE2 was used for co-incubation with 1nM COVID-QD treatments to compensate for the multiple Spike proteins conjugated to each construct. As a control, a 10nM soluble hACE2 treatment group was also used to ensure no basal stimulation or artefact may arise from the presence of exogenous hACE2. For URMC-099 rescue, a 200nM solution was prepared from a 100μM stock solution diluted in DMEM + 1% FBS. Prior to treatment with 10nM Spike protein or 1nM COVID-QD, the bEnd.3 monolayers were pre-treated with the URMC-099 solution for 1hr. URMC-099 was then aspirated off and replaced with treatment of 200nM URMC-099 + either 10nM Spike protein or 1nM COVID-QDs in DMEM + 1% FBS. A control treatment of just 200nM URMC-099 was also used to ensure no basal activity due to URMC-099. Rescue experiments involving either small molecule treatment was performed over 3 independent NVU co-cultures for each treatment. Prior to all treatments, the bEnd.3 monolayers were incubated in a reduced serum condition (DMEM + 1% FBS) for 3 hours.

#### Transendothelial Electrical Resistance (TEER)

An epithelial volt-ohm meter (World Precision Instruments; EVOM2) was used to take ensemble measurements of conductivity across endothelial cell contacts of bEnd.3 monolayers cultured on transwell inserts. TEER measurements were taken daily after bEnd.3 cultures appeared to have >80% confluency under a light microscope. A similar media composition of above and below the transwell membrane was used when taking TEER measurements and the media were allowed to equilibrate to RT for 30 minutes prior to measurements to reduce measurement artefacts due to temperature fluctuations. For bEnd.3 transwell cultures in co-culture with neuroglia, the inserts were measured in reduced serum media (DMEM + 1% FBS) prior to introduction of the transwell inserts into the co-culture. At the end of the treatment period, the transwell inserts were moved to a fresh 24-well culture plate containing DMEM + 1% FBS prior to taking a final TEER measurement. The reported TEER values are the difference between the treatment groups with the sham negative control group multiplied by the area of the transwell membrane.

#### Immunocytochemical Analysis

*Preparation of Immunolabeled Samples:* NVU co-cultures were first separated by removing the transwell inserts and placing them into a fresh 24-well plate with DMEM + 1% FBS. Both coverslips containing the primary neuroglia and the transwell inserts with the bEnd.3 monolayers were briefly washed with 1x Dulbecco’s phosphate buffered saline (DPBS, ThermoFisher; cat: 14190144), followed by fixation with 4% paraformaldehyde (PFA) in 1x DPBS for 15 minutes. This was followed by 5 minutes with 100mM glycine in 1x DPBS and a 5-minute washes with 1x DPBS. The cells were then permeabilized with 0.25% Triton-X (Millipore Sigma; cat: T9284) in 1x DPBS for 15 minutes, followed by two 5-minute washes with 1x DPBS. A 1-hour blocking step with 10% bovine serum albumin (BSA, Millipore Sigma; cat: A1470) in 1x DPBS was used after permeabilization and followed with treatment with the relevant primary antibodies (SI Appendix Table S1) in 3% BSA in 1x DPBS overnight on a rocker at 4^o^C. The next day, the cells were then washed for 5 minutes with 1x DPBS two times, followed by treatment with the secondary antibodies, as outlined in SI Appendix Table S2, in 3% BSA in 1x DPBS for 1 hour on a rocker at RT. The cells were then washed for 5 minutes with 0.1% Tween^®^ 20 (Millipore Sigma; cat: 655204) in 1x DPBS two times. The cells were then washed for 5 minutes in 1x DPBS. Coverslips were then dipped into ddH_2_O to remove any residual salt crystals and mounted on microscope slides (Fisher Scientific; cat: 22-034486) using ProLong^TM^ Diamond Antifade Mountant with DAPI (ThermoFisher; cat: P36962). The transwell membranes were cut out of the inserts before mounting onto microscope slides with an additional sealing layer with a rectangular #1.5 coverglass (Chemglass; cat: 48393-195) mounted on the membranes. The slides were allowed to cure overnight before imaging.

*Fluorescent Imaging w/ “Grid” Confocal Microscope:* The slides as prepared above were imaged on an Olympus BX51 microscope connected to a Hamatsu ORCA-ER detector and illuminated with a Prior Lumen 200 source with a Hg lamp (Prior; cat LM200B1-A). Excitation lines and bandpass emission filter (BPF) pairs are as follows – 350nm / DAPI (Semrock; cat: FF02-447/60-25); 405nm / FITC (Semrock; cat: FF01-524/24-25); 488nm / TRITC (Semrock; cat: FF01-593/40-25); 568nm / Cy5 (Semrock; cat: FF01-692/40-25); 350nm / TRITC – and were used to capture PL from DAPI, AlexaFluor488, AlexaFluor568, AlexaFluor647, and CdSe/CdS QD, respectively. The emission was collected through an infinity-corrected 20x UPlanApo 0.70 NA objective (Olympus). The emission is then passed through an OptiGrid structured illumination element to form a “grid” confocal image on the detector. For each sample, a z-stack was captured at interval steps of 1μm and compressed into the extended focus view presented in the representative images used in this manuscript. The exposure time for each channel was optimized and kept the same between each sample.

*Volocity Image Analysis:* The acquired z-stacks were analyzed using the Volocity 3D Image Analysis software (PerkinElmer). A fine noise filter was used on all images before applying a set of measurement protocols. For the bEnd.3 monolayers, a measurement protocol was designed to identify objects above a certain threshold corresponding to immunofluorescent labeled nuclei (λ_ex_ = 350nm, DAPI BPF), PECAM-1 (λ_ex_ = 405nm, FITC BPF), CLDN-5 (λ_ex_ = 488nm, TRITC BPF), and VCAM-1 (λ_ex_ = 568nm, Cy5 BPF), as well as CdSe/CdS QD emission (λ_ex_ = 350nm, TRITC BPF). The sum of the measured intensities and the sum of total spatial volume (i.e. voxels) from the objects were then exported for further analysis for the PECAM-1, CLDN-5, and VCAM-1 objects. The total number of detected nuclei and CdSe/CdS objects were also exported. For the neuroglial cultures, a measurement protocol was designed to identify objects above a certain threshold corresponding to immunofluorescent labeled nuclei (λ_ex_ = 350nm, DAPI BPF), PSD-95 (λ_ex_ = 405nm, FITC BPF), MAP2 (λ_ex_ = 488nm, TRITC BPF), and GFAP (λ_ex_ = 568nm, Cy5 BPF), as well as CdSe/CdS QD emission (λ_ex_ = 350nm, TRITC BPF). The sum of measured object intensities and spatial volume was extracted for the PSD-95, MAP-2, and GFAP objects. The MAP-2 objects were also further analyzed to extract the prevalence of dendritic beading by setting a cutoff for object volume and threshold for spheroidicity to identify true “beads.” The total number of beads, nuclei, and CdSe/CdS objects were also exported for further analysis.

#### Statistical Analysis

All quantitative values were organized and pre-processed on Excel prior to importing the values onto GraphPad Prism 9. Each replicate value was imported. For the rescue experiments of the NVU co-culture experiments, a two-way ANOVA with Holm-Sidak post-hoc correction was used. For all other experiments, a one-way ANOVA with Holm-Sidak post-hoc correction was used. Statistical significance was defined as an adjusted p-value less than 0.05 for all analyses.

### FIGURES

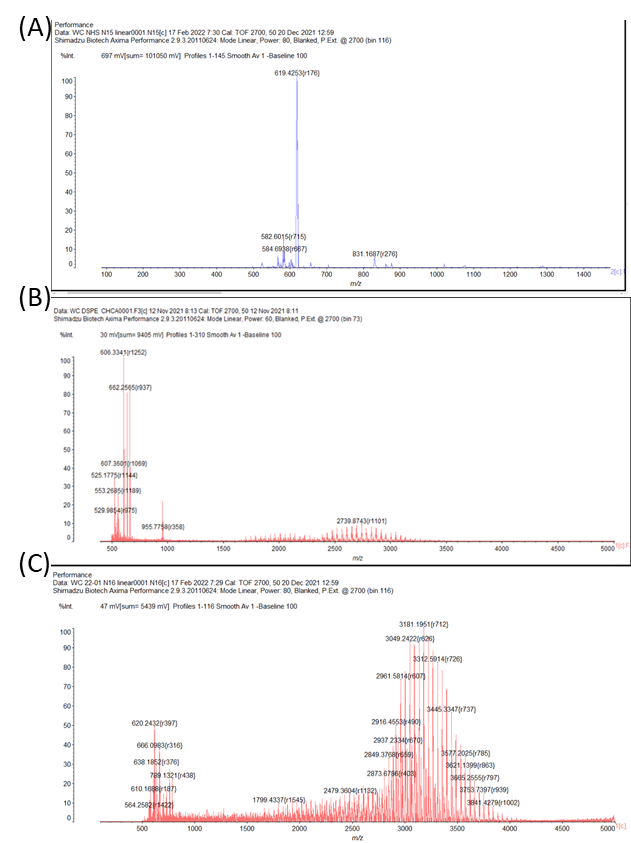

Figure S1. MALDI-ToF Spectra for (A) NHS-bis-sulfone (B) DSPE-PEG_2k_-NH_2_ and (C) PE:PEG:bis-sulfone.

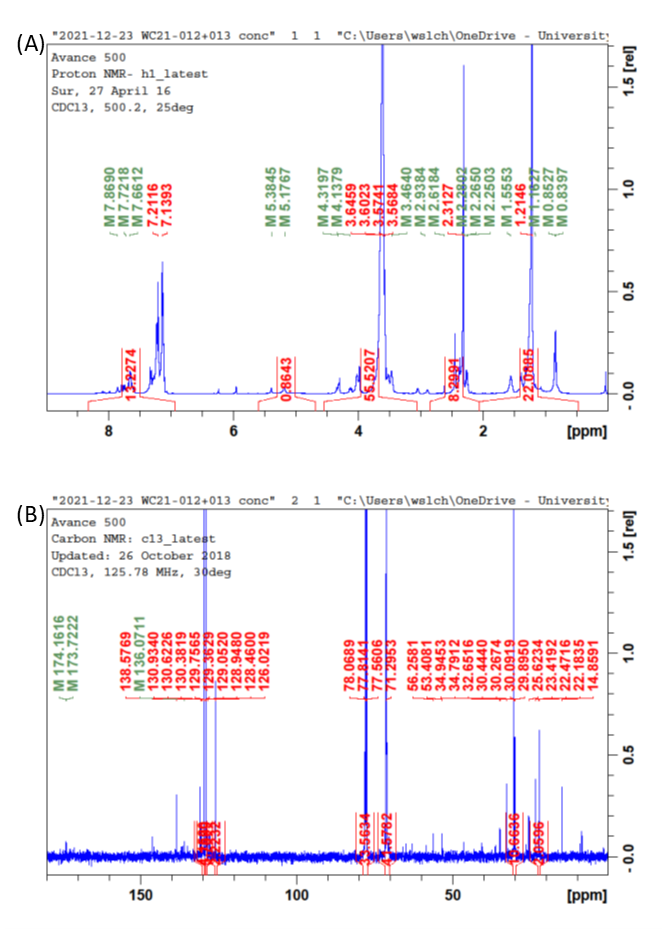

Figure S2. NMR spectra for PE:PEG:bis-sulfone (A) 1H (B) 13C.

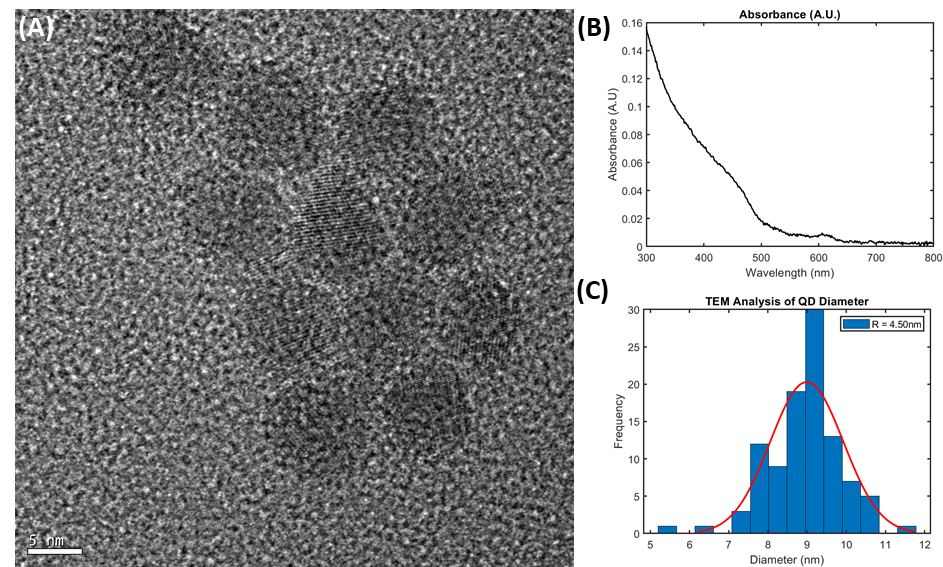

Figure S3. (A) Representative TEM image of CdSe/CdS QDs, scale bar = 5nm. (B) Absorbance of CdSe/CdS QDs. (C) Histogram of QD diameters taken from TEM images using ImageJ.

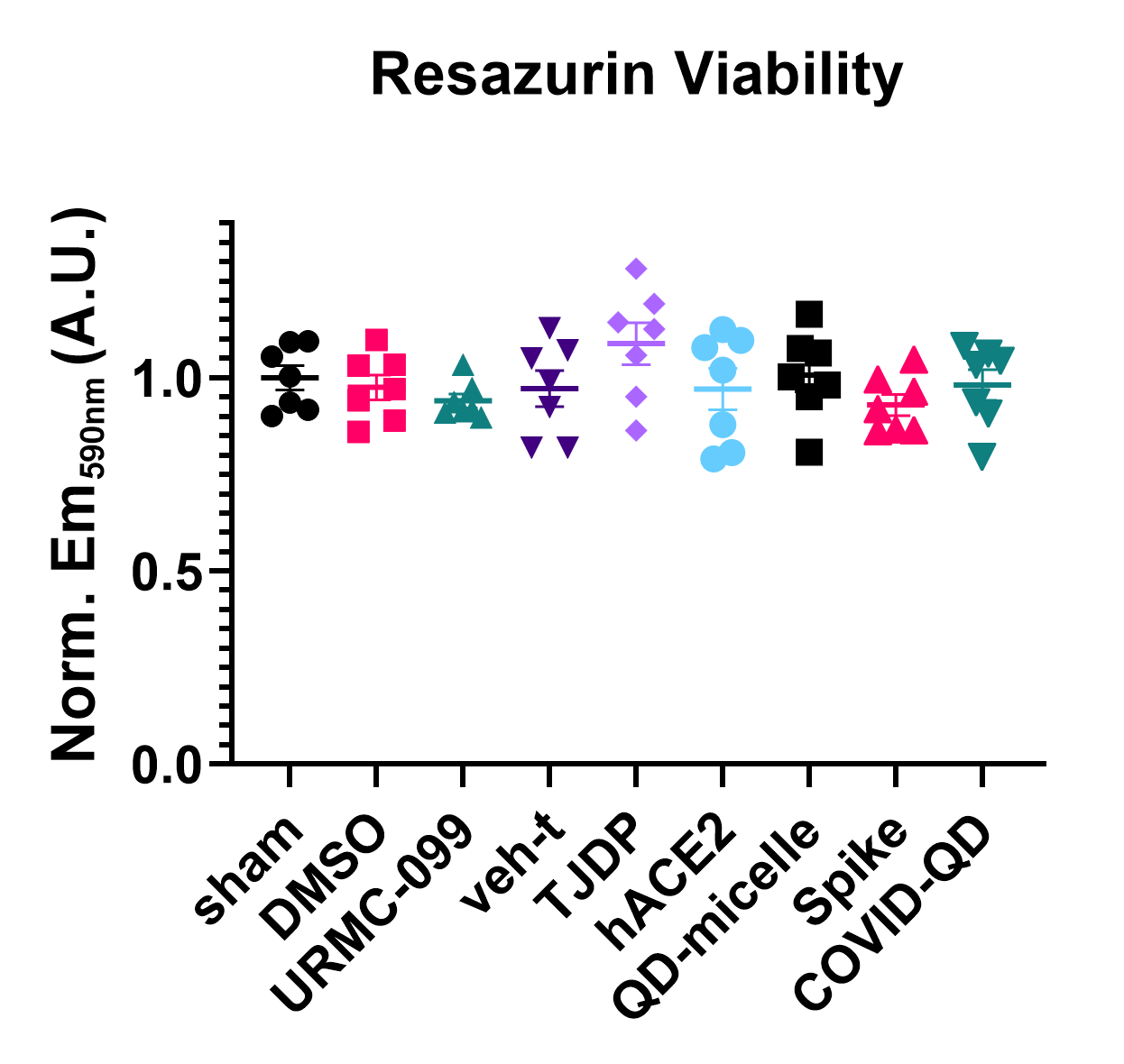

Figure S4. Resazurin metabolism viability assay of bEnd.3 cells in response to various independent reagents used in this study. Reported values are mean $\boldsymbol{\pm}$ SEM and all were found to be not significant (adjusted p $\boldsymbol{\geq}$ 0.05). n = 7 cultures pooled from two passages.

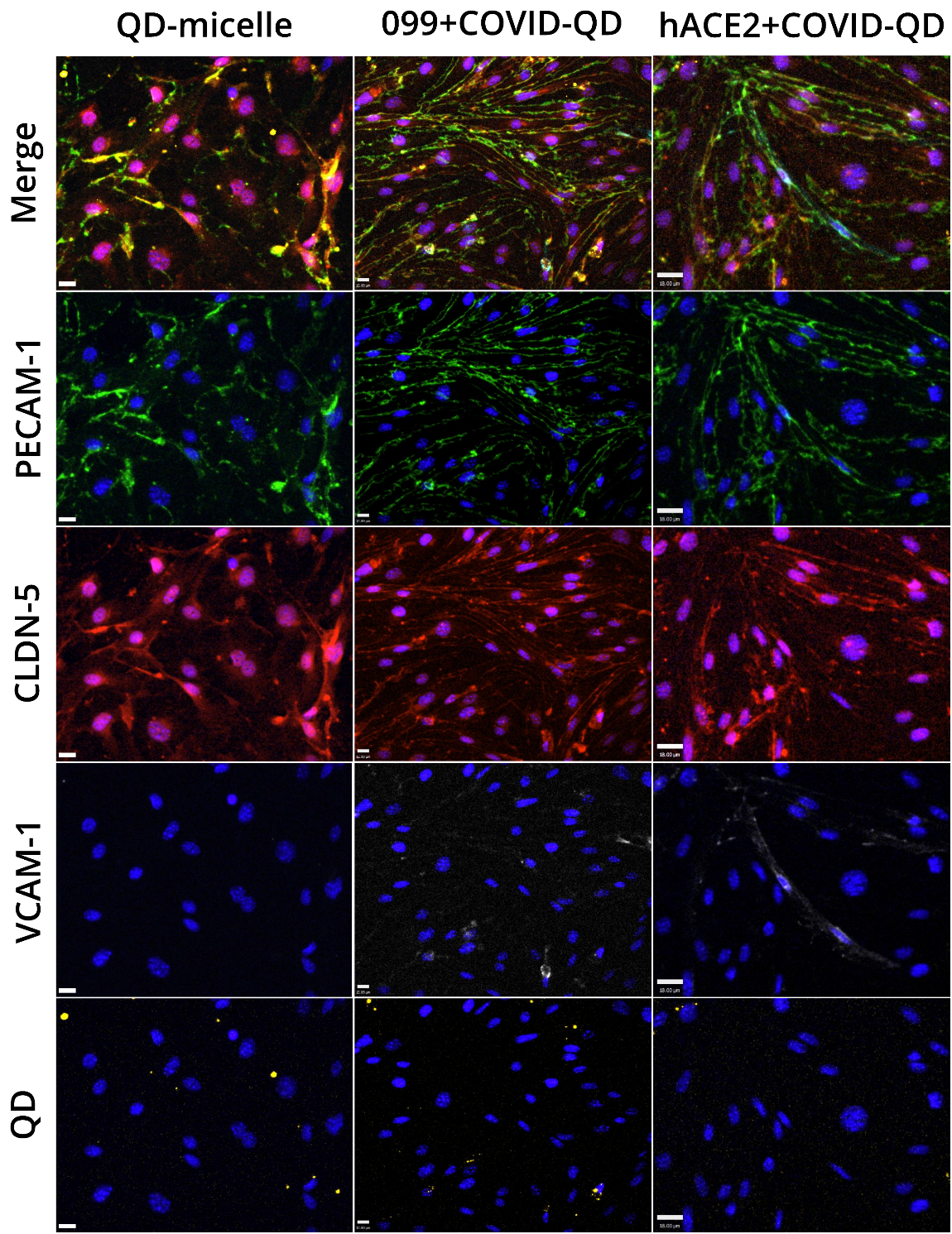

Figure S5. Immunofluorescent images of bEnd.3 monolayers treated with 1nM QD-micelles or 1nM COVID-QDs with either 200nM URMC-099 pre-treatment or 10nM soluble hACE2 co-treatment. Scale bars = 15μm. Nuclei stained by DAPI (deep blue) are shown in all panels.

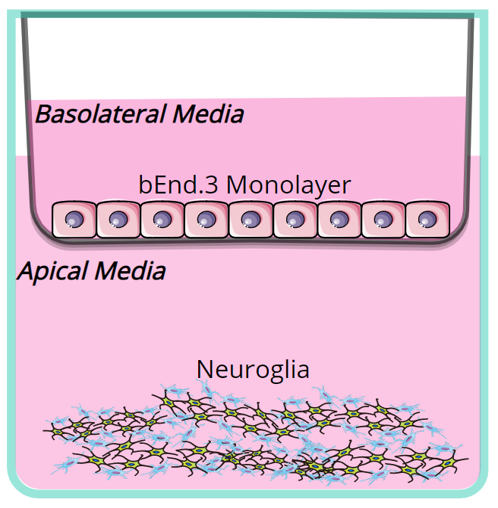

Figure S6. Setup of *in vitro* NVU co-culture model system.

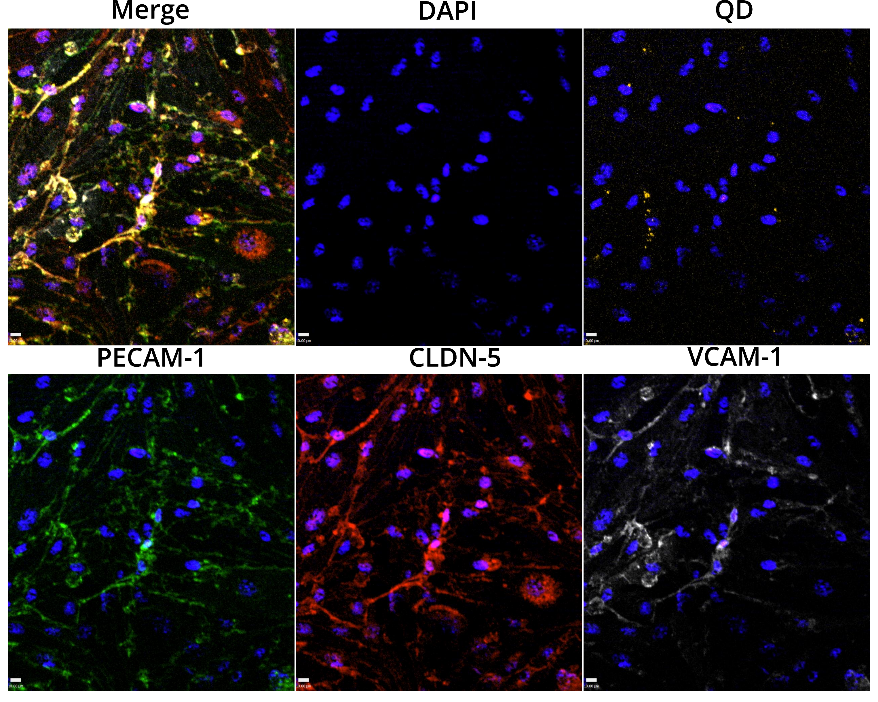

Figure S7. bEnd.3 monolayers treated with 1nM COVID-QDs exhibiting binding / presence of COVID-QDs on monolayer (yellow puncta). Other labels are PECAM-1 (green), CLDN-5 (red), VCAM-1 (white), nuclei (deep blue), and a merged channel. Scale bar = 15μm.

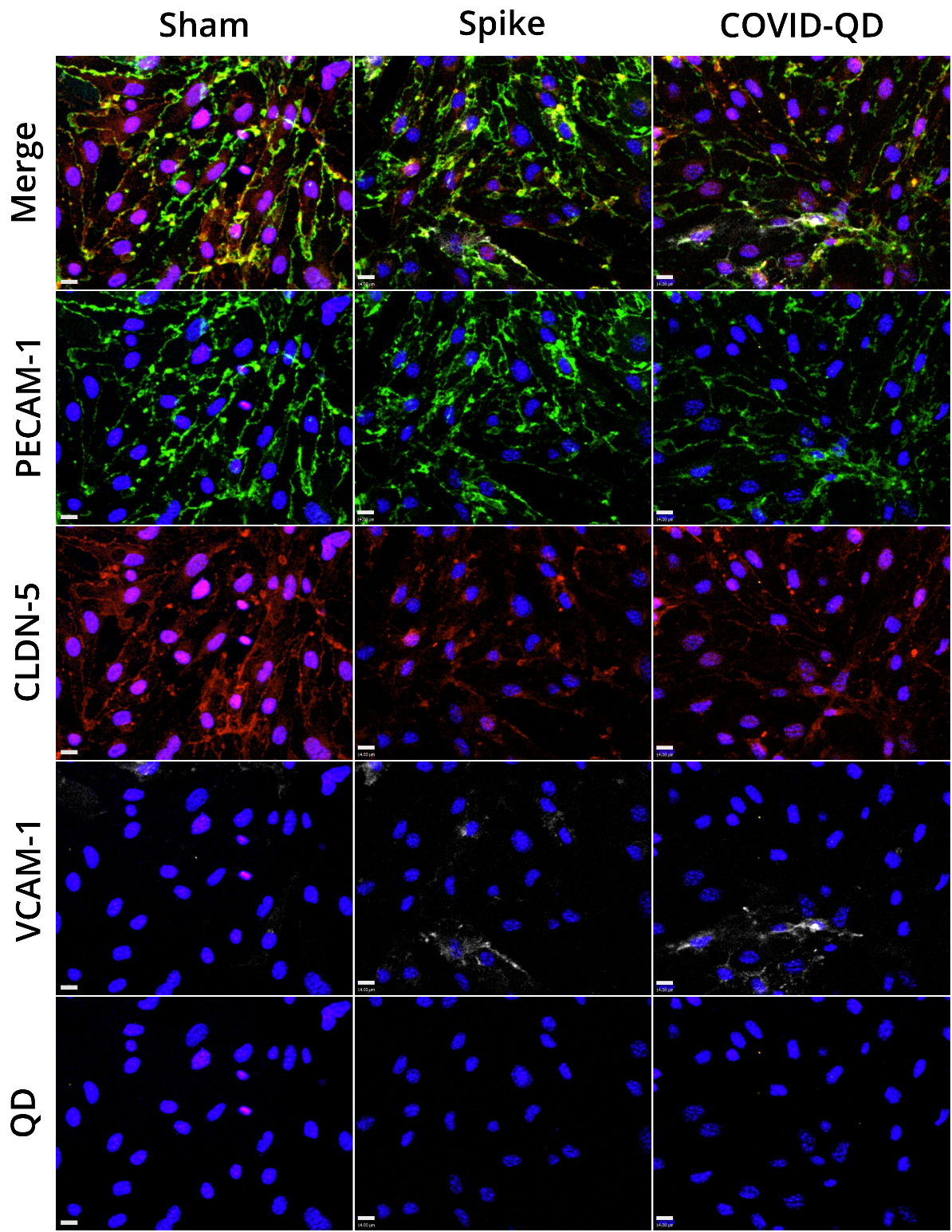

Figure S8. Dysregulation of bEnd.3 monolayer health in transwell co-cultures with neurons and astrocytes in response to either 10nM Spike protein or 1nM COVID-QDs compared to a sham no treatment group. Scale bars = 15μm. Nuclei stained by DAPI (deep blue) are shown in all panels.

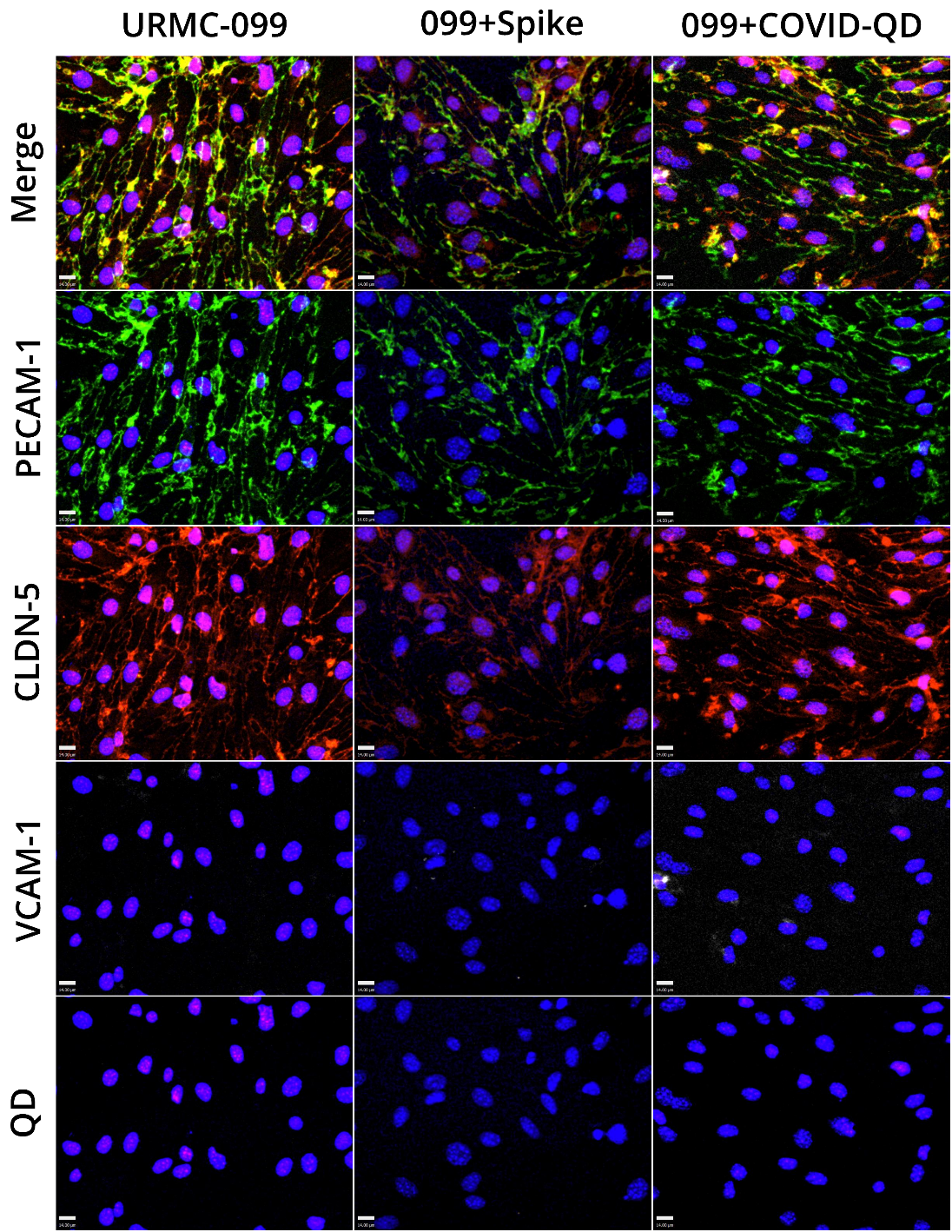

Figure S9. Rescue of bEnd.3 monolayers in transwell co-culture with neurons and astrocytes by pre-treating with 200nM URMC-099 before treatment with either 10nM of Spike protein or 1nM of COVID-QDs. Scale bars = 15μm. Nuclei stained by DAPI (deep blue) are shown in all panels.

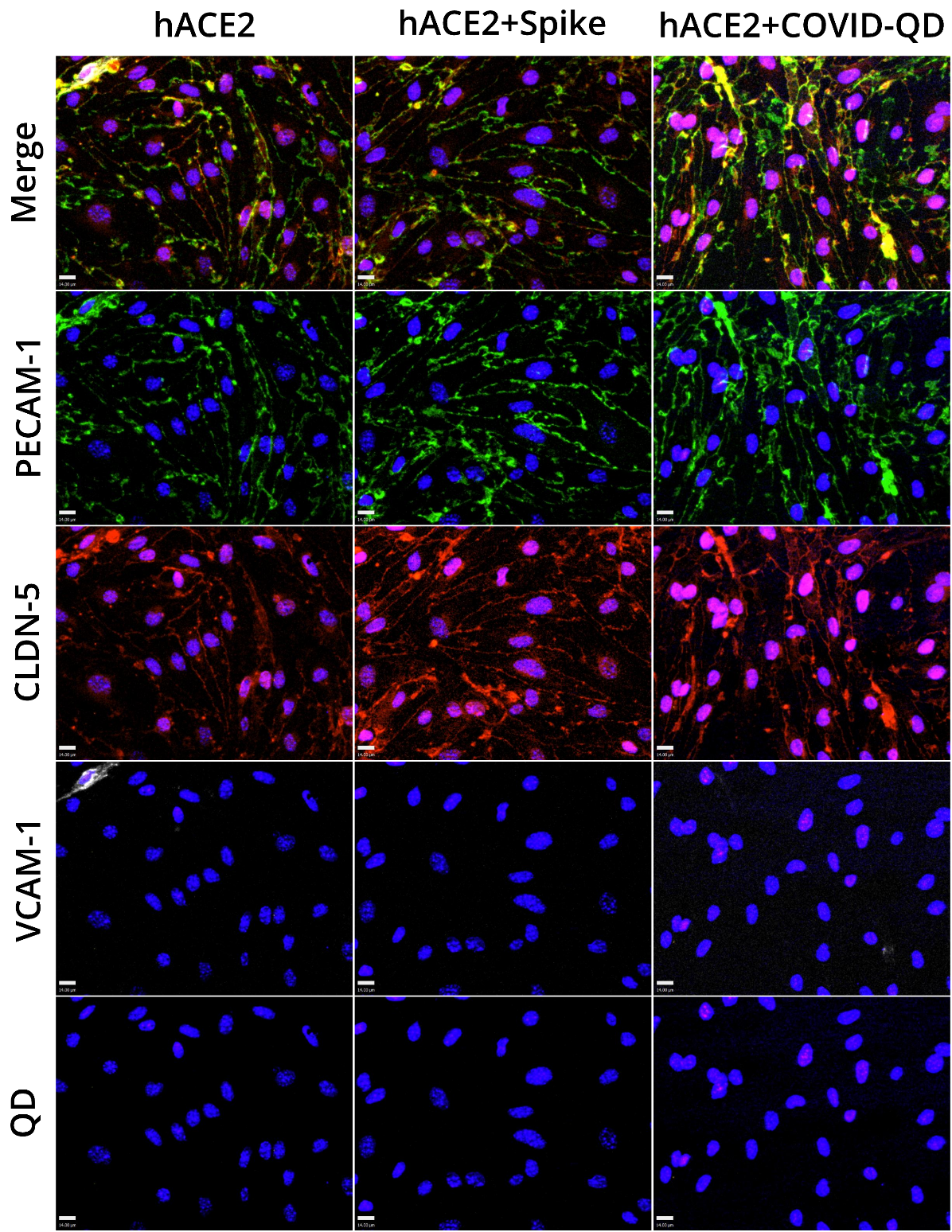

Figure S10. Rescue of bEnd.3 monolayers in transwell co-culture with neurons and astrocytes by co-treating 10nM of soluble hACE2 with either 10nM of Spike protein or 1nM of COVID-QDs. Scale bars = 15μm. Nuclei stained by DAPI (deep blue) are shown in all panels.

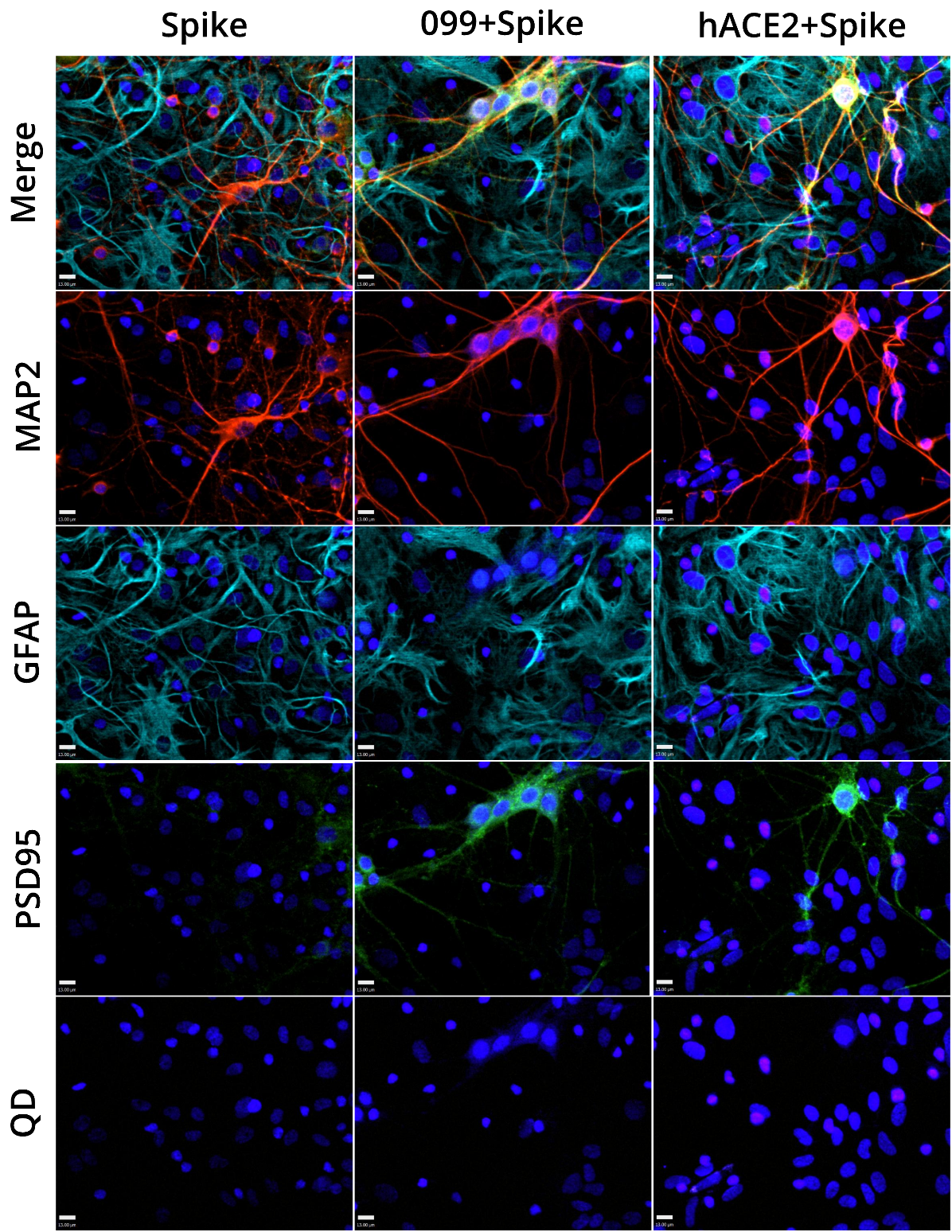

Figure S11. Small molecule rescue of neuron and astrocyte health by either 200nM URMC-099 pre-treatment or 10nM soluble hACE2 co-treatment with 10nM Spike protein. Scale bars = 15μm. Nuclei stained by DAPI (deep blue) are shown in all panels.

### TABLES

| Table S1. Selection of primary antibodies and dilutions used for immunofluorescent staining | | | | | |
| --- | --- | --- | --- | --- | --- |
| ***Culture*** | ***Host*** | ***Target*** | ***Dilution*** | ***Source*** | ***Catalog #*** |
| bEnd.3 | Hamster | PECAM-1 | 1:200 | ThermoFisher | MA3105 |
|  | Rabbit | CLDN-5 | 1:200 | ThermoFisher | PA5-99415 |
|  | Rat | VCAM-1 | 1:200 | BD Biosciences | 553330 |
| neurons | Mouse | PSD95 | 1:200 | NeuroMab | 75-028 |
|  | Rabbit | MAP2 | 1:150 | Cell Signaling | 4542 |
| astrocytes | Chicken | GFAP | 1:250 | Neuromics | CH22102 |

| Table S2. Selection of secondary antibodies used for immunofluorescent staining | | | | | |
| --- | --- | --- | --- | --- | --- |
| ***Host*** | ***Reactivity*** | ***Fluor*** | ***Dilution*** | ***Source*** | ***Catalog*** |
| Goat | Hamster | Alexa 488 | 1:750 | Jackson ImmunoResearch | 127-545-099 |
|  | Rabbit | Alexa 568 |  | ThermoFisher | A-11036 |
|  | Rat | Alexa 647 |  | ThermoFisher | A-21247 |
|  | Mouse | Alex 488 |  | ThermoFisher | A-11006 |
|  | Chicken | Alexa 647 |  | ThermoFisher | A-21449 |
